# ATM safeguards DNA replication at endogenous base lesions

**DOI:** 10.64898/2026.03.20.713105

**Authors:** Lucia Sommerova, Ashleigh King, J. Ross Chapman

**Affiliations:** Genome Integrity laboratory, Medical Research Council Molecular Haematology Unit, MRC Weatherall Institute of Molecular Medicine, Radcliffe Department of Medicine, The University of Oxford, Oxford, UK

## Abstract

Poly(ADP-ribose) polymerase inhibitors (PARPi) exploit homologous recombination deficiency (HRD) in BRCA1/2-mutated cancers to induce synthetic lethality^1–3^. PARPi also kill cancer cells lacking the DNA damage-responsive kinase ATM^4–6^, however, inconsistent evidence of HRD^7–9^ and variable clinical responses^9–11^ have obscured the underlying mechanism. Here we define how PARPi induce cytotoxicity in ATM-deficient cells and reveal a critical role for ATM in regulating DNA replication. In the absence of ATM, unrestrained PRIMPOL-dependent repriming at spontaneous oxidative base adducts generates discontinuous daughter strands containing DNA gaps that activate PARP. This defect is sustained by aberrant suppression of replication fork slowing – presumably via fork reversal – by the BRCA1-A complex, whose recruitment to stalled forks is normally counteracted by ATM. The resulting gaps require homologous recombination (HR) for post-replicative repair and underlie the synthetic lethal interaction with PARPi. Suppressing repriming, reducing oxidative stress, or blocking base excision repair alleviates these defects. Collectively, our findings reveal how spontaneous base damage cooperates with replication dysfunction to drive PARPi sensitivity and establish a paradigm of post-replicative repair addiction in ATM-deficient cells. Together, our findings define a mechanistic link between oxidative DNA damage and ATM-dependent replication control, illuminating how oxidative stress may exacerbate genome instability in Ataxia Telangiectasia.

## Main Text

Upon engaging DNA single-strand break (SSB) intermediates of base excision repair (BER) and/or lagging strand DNA synthesis, the activated PARP1/2 enzymes conjugate and polymerize long branched poly(ADP-ribose) (PAR) chains on themselves and the histones of nearby nucleosomes that play important functions in the signaling and repair of DNA damage^12,13^. Auto-PARylation of PARP1/2 promotes DNA repair primarily by recruiting the PAR-binding protein XRCC1, the central scaffolding component of the SSB repair pathway that recruits the repair enzymes including DNA ligase III (LIG3) and DNA polymerase β (POLB)^14^. PARP1/2 autocatalysis also enables the release of activated PARP1/2 proteins from SSBs^12,15^. Thus, by blocking PARP1/2 autocatalysis, small molecule catalytic inhibitors of PARP1/2 (PARPi) lead to PARP1/2-bound DNA complexes that are physical barriers to DNA replication and repair^15^, in addition to blocking SSB repair protein mobilization^14^. During DNA replication, replication fork convergence with PARPi-induced PARP-DNA complexes results in DNA repair intermediates including DSBs whose repair requires homologous recombination (HR). Consequently, tumors deficient in HR, such as those harboring *BRCA1* or *BRCA2* mutations, are exquisitely sensitive to PARPi treatment^1,2^. Indeed, the selectivity of PARPi towards inducing synthetic lethality in HRD cells, while sparing normal tissues, has underpinned the successful use of PARPi in BRCA1/2-mutated breast, ovarian, and prostate cancers.

The therapeutic success of PARPi in *BRCA*-mutated tumors has motivated their evaluation in additional DNA repair-deficient cancers, including those with pathogenic alterations in the *Ataxia Telangiectasia Mutated (ATM)* gene. The ATM kinase is a master regulator of the cellular response to DNA double-strand breaks (DSBs), initiating DNA damage checkpoint signaling, coordinating DNA repair, and modulating oxidative stress responses through phosphorylation of numerous substrates involved in redox homeostasis and metabolism^16–18^. Germline biallelic *ATM* mutations cause Ataxia Telangiectasia (A-T), a neurodegenerative and cancer-predisposition syndrome^19^, while heterozygous carriers display increased susceptibility to breast and other cancers^20^. Somatic *ATM* alterations are also frequent in prostate cancer (PCa), where they associate with aggressive disease^21^.

Early studies showed that depletion or pharmacological inhibition of ATM hypersensitizes cancer cells to PARPi-induced cytotoxicity^4–6^, signifying the potential of PARPi therapy in ATM-deficient tumors^3^. However, clinical trials in ATM-mutated prostate cancer have yielded inconsistent responses^9–11^. Furthermore, while initial work attributed PARPi sensitivity in *ATM-*deficient cancer cells to HRD^4,5^, subsequent studies have challenged this model. Reporter assays showed HR capacity to be largely intact in *Atm*-knockout mouse fibroblasts and embryonic stem cells^7,22,23^. Likewise, PARPi-induced recruitment of the HR recombinase RAD51 was normal in *ATM*-deficient PCa cells^9^. Thus, the DNA damage response defect driving PARPi cytotoxicity in *ATM-*deficient cells, the reasons for its variable presentation in a clinical setting, and its pathophysiological significance for A-T remain unresolved problems of medical significance.

In this work, we define how ATM safeguards DNA replication at endogenous oxidative base lesions, establishing a unifying mechanism that links replication dysfunction, oxidative DNA damage, and cellular sensitivity to PARP inhibition.

### *ATM*-deficient cells accumulate replication-associated single-strand DNA breaks

The hypersensitivity of *BRCA* mutated tumors to both treatments with platinum salts and PARPi is a hallmark of HRD that is widely exploited in clinical practice^8^. The role of HR in fostering cross-resistance to both PARPi and cisplatin can be clearly demonstrated using a *BARD1^AID/AID^* HCT-116 cell line in which BRCA1 can be conditionally inactivated upon auxin-induced degradation of its obligate binding partner BARD1 (Figure 1a,b; hereafter referred to as *BARD1*^Δ*/*Δ^ cells)^24^. We were therefore intrigued when we found that *ATM-*deleted HCT-116 cells, by contrast, exhibited olaparib hypersensitivity, yet retained wild type levels of cisplatin resistance in parallel experiments (Figure 1a,b). To question the basis of these differences, we quantified RAD51 recruitment into nuclear foci, as a proxy for HR activity in control (*BARD1^AID/AID^* HCT-116 untreated with auxin), *ATM^−/−^* and *BARD1*^Δ*/*Δ^ HCT-116 cells treated with olaparib for 24 h, and Synthesis (S)-phase staged cells 2 h following ionizing radiation exposure. In these experiments, *ATM^−/−^* cells showed wild type levels of RAD51 foci following either treatment (Figure 1c and Figure S1a,b). By contrast, the frequencies of RAD51 foci were severely diminished in *BARD1*^Δ*/*Δ^ cells, consistent with their defect in HR. Consistent results were obtained when we quantified spontaneous sister chromatid exchange events (SCEs) in metaphase chromosome spreads as a proxy for HR-dependent recombination frequencies in wild type, *ATM^−/−^* and *BARD1*^Δ*/*Δ^ cells. SCEs occurred at comparable frequencies in wild type and *ATM^−/−^* cells yet were significantly decreased in HR-deficient *BARD1*^Δ*/*Δ^ cells, as would be expected (Figure S1c,d).

**Figure 1:**
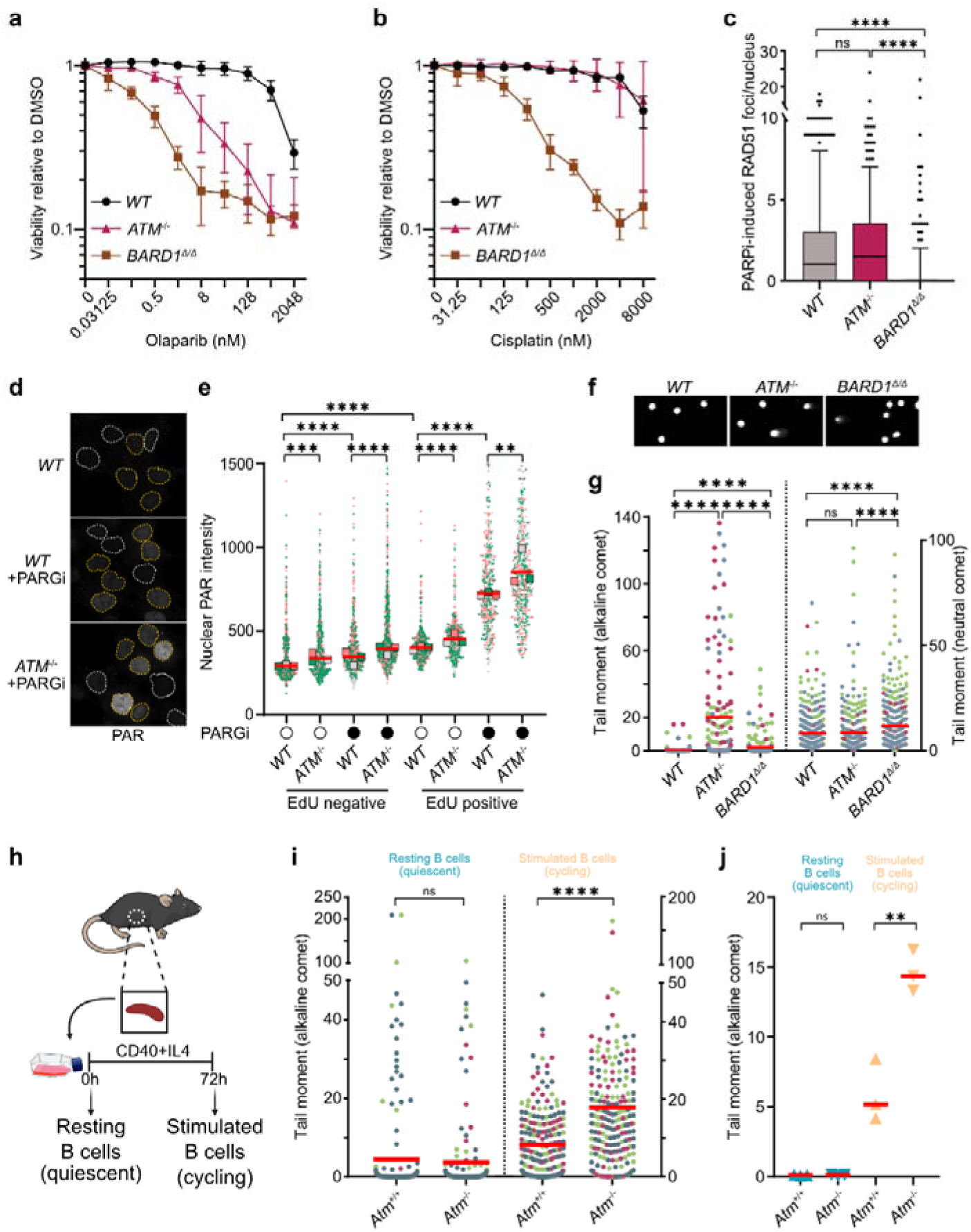
PARP-activating SSBs accumulate in replicating *ATM*-deficient cells. **a,b,** Survival of the indicated *BARD1^AID/AID^* HCT-116 cell lines exposed to the indicated doses of olaparib (**a**) or cisplatin (**b**). Resazurin cell viability assay, *n=3* biological experiments, mean ± s.d. **c,** Immunofluorescence imaging-based quantification of RAD51 foci in indicated *BARD1^AID/AID^* HCT-116 cell lines following 24 h treatments with olaparib (*n=2* biological experiments, >200 cells per condition). Horizontal bars, boxes and whiskers denote median, 25–75, and 5–95 percentile values, respectively. Significance, two-sided Kruskal–Wallis H test with Dunn’s correction for multiple comparisons. ****P ≤ 0.0001; ns non-significant. **d,e,** Immunofluorescence imaging of PAR levels in indicated HCT-116 cells treated with 1 mM PARGi or mock (DMSO) for 45 min and pulse-labelled with EdU (20 min, 10 µM). Representative cells (**d**), EdU -positive and -negative nuclei are outlined in yellow and white, respectively. Superplots (**e**), integrated PAR intensity per nucleus, *n=3* biological experiments (each color denotes one replicate, bars indicate median and colored squares indicate median PAR signal per replicate). Significance, two-way ANOVA mixed model with repeat measures and Sidak’s correction for multiple comparisons. ****P ≤ 0.0001, ***P ≤ 0.001 **P ≤ 0.01. **f,g** DNA damage quantification in the indicated *BARD1^AID/AID^* HCT-116 cells by comet assay under alkaline or neutral conditions. Representative images (**f**), and Scatter plots (**g**), show comet tail moments across *n=3* biological experiments, each comprising 80-100 cells and represented in a different color. Horizontal bars, median values. Significance, two sided Kruskal-Wallis H test with Dunn’s correction for multiple comparisons. ****P ≤ 0.0001. **h,** Experimental design for **i,j** using splenic B cells. **i,j** Alkaline comet tail moments in quiescent resting and stimulated cycling cultures of splenic B cells from indicated mice. Scatter plot (**i**), each color represents one of n = 3 biological experiments; horizontal bars denote medians. Significance, Mann-Whitney t-test, ****P ≤ 0.0001; ns, non-significant. Scatter plot (**j**), Median tail moment in *n=3* biological experiments, with indicated mice. Significance, unpaired two-tailed t-test, **P ≤ 0.01; ns, non-significant. Throughout the figure, *WT* denotes non-auxin-treated *BARD1^AID/AID^* cells. For *BARD1*^Δ*/*Δ^ cells, *BARD1^AID/AID^*cell lines were treated with doxycycline (2 μg/ml, 24 h) before addition of auxin (IAA, 1 mM) or dimethyl sulfoxide (DMSO) (carrier control), followed by olaparib.

The insensitivity of *ATM*^−/−^ cells to cisplatin, combined with their positive indicators of HR activity, prompted us to consider an alternative explanation for the PARPi hypersensitivity of *ATM*-deficient cells. We therefore considered that *ATM-*deficient cells might spontaneously accumulate DNA damage that activates PARPs to promote DNA repair. To test this, quantitative immunofluorescence microscopy was used to measure PAR levels in the nuclei of wild type, *ATM^−/−^,* and *BARD1*^Δ*/*Δ^ HCT-116 cells. PARP-dependent signaling is highest during S phase, mostly due to ssDNA exposure at sites of persistent unligated Okazaki fragments during lagging strand DNA synthesis^25,26^. Thus, we used EdU pulse-labelling to discriminate S phase and G1/G2 -staged cells, in experiments where the cells were also either mock-treated, or exposed to an inhibitor of the poly(ADP-ribose) glycohydrolase (PARGi; PDD00017273) to stabilize accumulated nuclear PAR. Consistent with prior reports^25,26^, PAR-signals were lowest in non-S phase (EdU-negative) nuclei, where they only slightly increased upon exposure to PARGi (Figure 1d,e and Figure S2a). By contrast, nuclear PAR was elevated in S phase staged nuclei in all cell lines (Figure 1d,e and Figure S2a). Interestingly, nuclear PAR levels were significantly higher in *ATM^−/−^* nuclei, than they were in wild type and *BARD1*^Δ*/*Δ^ cells, with these differences further exacerbated upon PAR stabilization with PARGi treatment (Figure 1d,e and Figure S2a). Catalytic inhibition of ATM with the ATM inhibitor (ATMi) AZD0156 also reproducibly increased PAR signals in wild type HCT-116, the non-cancerous cell line hTERT RPE-1, and the triple negative breast cancer cell line MDA-MB-231 (Figure S2b-d).

Elevated PARP signaling in *ATM* deleted or inhibited cells was suggestive of spontaneous DNA damage accumulation, leading us to investigate what lesions were activating PARP in *ATM*-deficient cells. To this end, we conducted comet assays with wild type, *ATM^−/−^,* and *BARD1*^Δ*/*Δ^ HCT-116 cells, where subjection of agarose-embedded cells to electrophoresis under alkaline-denatured or neutral conditions, enables for the resolution of SSBs, alkali-labile sites and DSBs, or DSBs, respectively. ∼40% of *ATM^−/−^* nuclei resolved in alkaline comet assays exhibited tail moments indicative of unrepaired DNA strand breaks and lesions, that were notably absent from the majority of nuclei from control or HR-deficient *BARD1*^Δ*/*Δ^ cells (Figure 1f,g and Figure S2e). However, when performed under neutral conditions, the tail moment signals in wild type and *ATM^−/−^* nuclei were equivalent, confirming that *ATM*-deficient cells do not accumulate higher levels of spontaneous DSBs (Figure 1g). This implies that the alkaline tail moments in *ATM^−/−^* nuclei are primarily caused by SSBs. Of note, these results clearly contrasted with *BARD1*^Δ*/*Δ^ cells, where significant increases in neutral comet tail moments indicated the presence of spontaneously occurring DSBs (Figure 1g). This increased DSB burden in *BARD1-*deficient cells likely accounts for the modest increase in tail moments also seen in alkaline comet assays. Importantly, equivalent *ATM*-knockout associated increases of SSB signal (without increases in DSBs) were observed when comet assays were replicated with wild type and *ATM-*knockout clones of the hTERT RPE-1 cell line (Figure S2f). Taken together, our results reveal an ATM-dependent suppression of SSB accumulation in cultured human cells, substantiating earlier reports of elevated spontaneous SSBs in ATM-deficient settings^27,28^.

PAR accumulation was only evident in S phase staged *ATM^−/−^*cells (Figure 1d,e), a fraction that correlated to the ∼40% of *ATM^−/−^* nuclei that were positive for tail moments in alkaline comet assays (Figure S2e). Thus, we next investigated the possible link between SSB formation and DNA replication, using primary B cells isolated from the spleens of wild type and *Atm-*knockout mice, contrasting SSB signals from freshly isolated quiescent B cells to fast-cycling B cells stimulated *ex vivo* (Figure 1h). Indeed, while nuclei from resting B cells isolates from either wild type or *Atm^−/−^* mice showed equivalent SSB signals, in stimulated B splenocytes, *Atm^−/−^* cell nuclei yielded increased tail moment signals that were ∼3-fold higher than that of wild type nuclei (Figure 1i,j). Lastly, we found that SSB signals in *ATM*-knockout HCT-116 cells could be quenched by inhibiting DNA synthesis with the polymerase poison aphidicolin yet were unaffected upon transcription inhibition using the RNA Pol2 inhibitor DRB (Figure S2g-i). Altogether, these results confirmed DNA replication as the driver of SSB formations in *ATM-*knockout cells, thus implicating ATM in suppressing replication-associated DNA damage.

### PARPi toxicity is a consequence of aberrant replication repriming in *ATM*-deficient cells

Discontinuity within the nascent strands synthesized during DNA replication represents a natural source of PARP-activating SSBs, with unligated Okazaki fragments representing the major source of these lesions^25,26^. In agreement, RPE-1 cells deleted of flap-endonuclease 1 (FEN1), an enzyme that mediates Okazaki-fragment trimming prior to ligation, displayed dramatic increases in nuclear PAR (Figure S2j). Elevated PAR signals in S-phase fractions of ATMi-treated cells by comparison were modest, rendering a role for ATM in conventional Okazaki fragment maturation unlikely. Nevertheless, to assess whether replication-dependent nascent strand gaps contribute to the PARP-hyperactivity seen in *ATM-*deficient cells, we used a modified version of DNA fiber analysis, in which single-strand nicks and gaps in nascent DNA strands renders IdU-labelled fibers susceptible to S1 nuclease cleavage (Figure 2a). In these experiments, the length of replicative fibers prepared from wild type cells was unaffected by S1 nuclease treatments (Figure 2b), signifying an absence of ssDNA gaps from most nascent DNA strands. By contrast, IdU tracts in fibers prepared from *ATM^−/−^* cells were consistently shortened following S1 nuclease treatments, confirming an abundance of ssDNA gaps in the nascent replicated strands (Figure 2b). In consideration of replication repriming, potentially at stalled replication forks, as a cause of post-replicative ssDNA gaps in *ATM*-deficient cells, we simultaneously measured the S1 sensitivity of replication fibers from ATM/PRIMPOL co-deleted cells. PRIMPOL is a bi-functional primase-polymerase enzyme that can enable leading strand repriming at modified/damaged bases^29^. Remarkably, S1 nuclease dependent replication fiber shortening was lost in *ATM^−/−^PRIMPOL^−/−^* cells (Figure 2b). PRIMPOL-deletion also blocked SSB accumulation in *ATM^−/−^PRIMPOL^−/−^* cells, as exemplified in alkaline comet assays relative to the parental *ATM^−/−^* cell line (Figure 2c). Finally, deletion of PRIMPOL in *ATM^−/−^* cells led to a complete loss of olaparib sensitivity (Figure 2d). Considered together, this data reveals that unrestrained replication repriming by PRIMPOL characterizes *ATM*-deficient cells, driving the accumulation of nascent-strand ssDNA gaps. Moreover, since these lesions stimulate PARP activity during replication, they reconcile why *ATM*-deficient cells are hypersensitive to PARPi despite retaining a normal capacity for HR.

**Figure 2:**
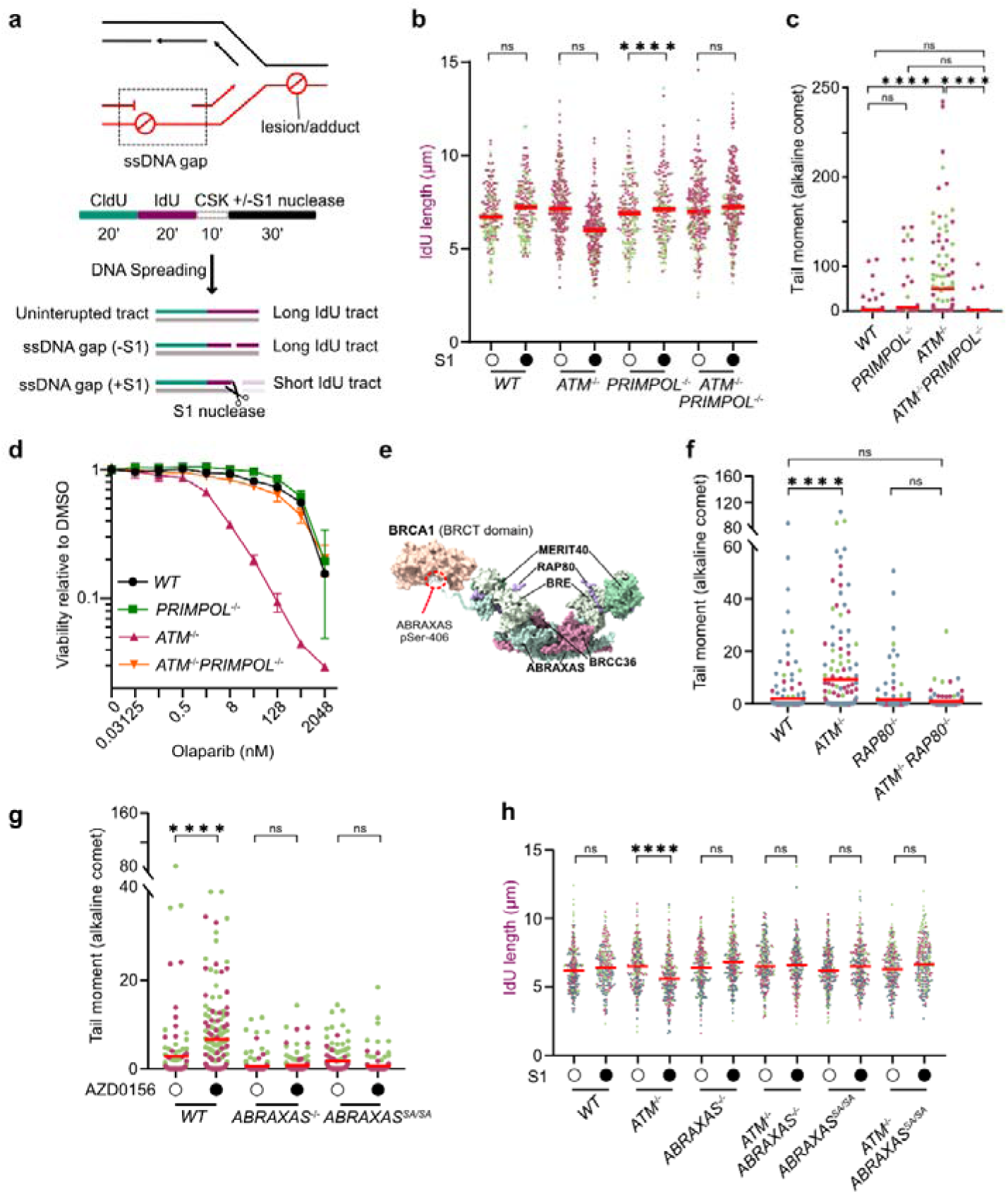
Replication repriming drives PARPi sensitivity in *ATM*-deficient cells. **a,** Schematic of nascent-strand SSB detection by S1-nuclease coupled DNA fiber assay. IdU, iododeoxyuridine; CldU, chlorodeoxyuridine; CSK, cytoskeletal buffer. **b,** Scatter plots showing IdU tract lengths in DNA fibers from the indicated *BARD1^AID/AID^* HCT-116 cells lines. Horizontal bars, median values of >250 fibers per cell line in *n=2* biological experiments, each represented in a different color. **c,** Alkaline comet tail moment data, *n=2* biological experiments (80–100 cells per experiment; each color denotes one replicate). Horizontal bars indicate medians. **d,** Survival assay of indicated *BARD1^AID/AID^* HCT-116 cell lines measured by resazurin assay 7 days following olaparib treatment. Data, represent mean ± s.d. from *n=3* biological experiments. **e,** Schematic representing the structure and function of BRCA1-A complex. **f,g** Alkaline comet tail moment in indicated cell lines. Data from *n=3* **(f)** or *n=2* **(g)** biological experiments (80–100 cells per experiment; each color denotes one replicate), Horizontal bars, median. **h,**Scatter plots, showing IdU tract lengths in S1 nuclease-coupled DNA fiber assay from the indicated *BARD1^AID/AID^* HCT-116 cells lines. Horizontal bars, median values of >250 fibers per cell line in *n=3* biological experiments, each represented in a different color. Throughout the figure, *WT* refers to non-auxin-treated *BARD1^AID/AID^* cells and significance was determined by two sided Kruskal-Wallis H test with Dunn’s correction for multiple comparisons. ****P ≤ 0.0001; ns, non-significant.

### The BRCA1-A complex sustains replication repriming in *ATM*-deficient cells

We next asked how ATM suppresses excessive repriming events. A recent mass spectrometry-based analysis of nascent chromatin identified ATM as selectively enriched at camptothecin (CPT)-stalled replication forks, where its kinase activity suppressed aberrant accumulation of DNA repair proteins^30^. Among these were all subunits of the BRCA1-A complex - a BRCA1-binding heteropentameric complex of two copies each of RAP80, BRE, MERIT40, BRCC36, and ABRAXAS (Figure 2e)^31^. Given that BRCA1-A complex has been linked to the CPT and PARPi hypersensitivity in *ATM*-deficient cells^32^, and to ATMi-induced cytotoxicity in cancer cells^33^, we examined whether abnormal BRCA1-A recruitment to replication sites contributes to replication defects in *ATM*-deficient cells. Alkaline comet assay revealed that steady-state SSBs in *ATM*-knockout cells were completely suppressed by co-deletion of *RAP80* (Figure 2f). Likewise, treatment with ATMi induced SSB accumulation in wild type cells, but not in *RAP80^−/−^* cells (Figure S3a). In both cases, DNA damage suppression correlated with a substantial, though incomplete, reduction in PARPi hypersensitivity (Figure S3b,c). Altogether, these results implicated BRCA1-A in the generation of PARP-activating SSBs in *ATM*-deficient cells, reconciling its reported contribution to PARPi sensitivity in this context^32^.

To dissect the mechanism underlying BRCA1-A-dependent damage, we examined the roles of its subunits. RAP80 binds K63-linked ubiquitin chains via its tandem ubiquitin-interacting motifs (UIMs 1/2), and BRCC36 provides K63-specific deubiquitinase (DUB) activity (Figure 2e). However, wild-type RAP80 and UIM1/2 mutated RAP80^AA,AA-RR,RR^ mutant protein equally restored alkaline comet signals in *ATM^−/−^ RAP80^−/−^* cells, despite the latter being unable to localize to DSB sites in irradiated cells (Figure S4a,b), as expected^34^. Similarly, ATMi induced SSBs accumulation in wild type cells and in cells expressing a DUB-inactive *BRCC36^QSQ/QSQ^* alleles^35^, but not in *BRCC36*-knockout cells (Figure S4c,d). These results indicate that BRCA1-A’s structural rather than catalytic properties promote SSB generation.

However, ATMi-induced DNA damage was abolished in *ABRAXAS*-knockout and *ABRAXAS^S^*^406^*^A/S^*^406^*^A^* cells where the BRCA1-binding phospho-epitope at Ser-406 is mutated (Figure 2g and Figure S4c), consistent with a role for BRCA1 interplay with the BRCA1-A complex in this context. Concordantly, PARPi hypersensitivity was reduced in *ABRAXAS^−/−^*, *ABRAXAS^S^*^406^*^A/S^*^406^*^A^*, *RAP80^−/−^*, and *BRCC36^−/−^* cells, but not in cells with DUB-inactive BRCC36 or UIM mutant RAP80 (Figure S4e-i). Finally, DNA fiber assays furthermore confirmed a role for ABRAXAS and its interaction with BRCA1 in the generation of nascent-strand ssDNA gaps in ATM-deficient cells (Figure 2h). Collectively, these results demonstrate that a structurally intact, BRCA1-bound BRCA1-A complex drives the generation of post-replicative ssDNA gaps in *ATM*-deficient cells.

### BRCA1-A supports replication-stress resistant DNA synthesis in *ATM-*deficient cells

Mammalian cells employ replication fork reversal to pause DNA synthesis at replication-blocking lesions allowing lesion resolution before resumption of DNA synthesis^36–38^. As replication fork reversal also counteracts replication repriming by PRIMPOL^39,40^, we hypothesized that the unrestrained replication repriming observed in *ATM*-deficient cells reflects defects in replication fork reversal (Figure 3a).

**Figure 3:**
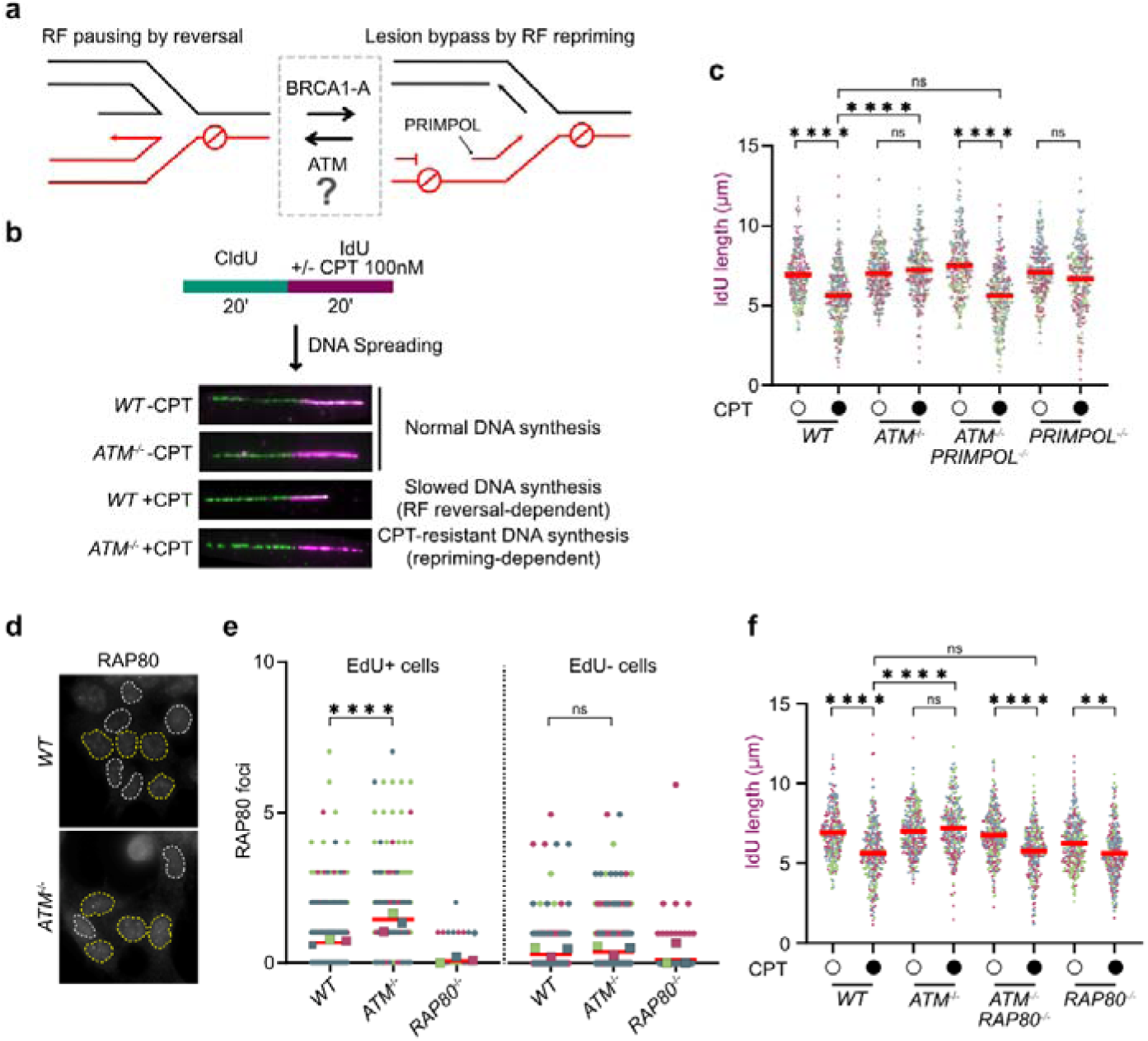
BRCA1-A supports replication-stress resistant DNA synthesis in *ATM*-deficient cells. **a,** Model illustrating the proposed roles of ATM and BRCA1-A complex at stalled replication fork, balancing replication fork reversal and repriming. **b,** Schematic of DNA fiber measuring CPT-induced replication slowing. IdU, iododeoxyuridine; CldU, chlorodeoxyuridine; CPT, campothecin. **c,** Quantification of DNA fiber tract lengths from CPT-treated indicated *BARD1^AID/AID^* HCT-116 cells lines. Horizontal bars, median values of >250 fibers per condition across *n=3* biological experiments, each represented in a different color. Significance, two sided Kruskal-Wallis H test with Dunn’s correction for multiple comparisons. ****P ≤ 0.0001; ns, non-significant. **d,e,** Immunofluorescence imaging of spontaneous RAP80 foci levels in indicated HCT-116 cell lines, pulse-labelled with EdU (5 min, 40 µM). Representative images (**d**), EdU -positive and -negative nuclei are outlined in yellow and white, respectively. Superplots **(e)** representing RAP80 foci per EdU positive and negative nucleus from *n=3* biological replicates (each color denotes one replicate; bars indicate means; colored squares, mean RAP80 foci per replicate). Significance, two-way ANOVA mixed model with repeat measures and Sidak’s correction for multiple comparisons. ****P ≤ 0.0001; ns, non-significant. **f,** Quantification of DNA fiber analysis from CPT-treated indicated *BARD1^AID/AID^* HCT-116 cells lines. Horizontal bars, median values of >250 fibers per condition across *n=3* biological experiments, each represented in a different color. Significance, two sided Kruskal-Wallis H test with Dunn’s correction for multiple comparisons. ****P ≤ 0.0001; ns, non-significant. Throughout the figure, *WT* refers to non-auxin-treated *BARD1^AID/AID^* cells.

To test this, we used DNA fiber analysis to monitor replication stress-induced replication fork slowing, as a recognized proxy for replication fork reversal^41^. In wild type cells, low dose of topoisomerase-1 (TOP1) inhibitor camptothecin (CPT) shortened IdU track lengths (Figure 3b,c), consistent with our expectation of fork reversal at sites of trapped TOP1-DNA lesions^42^. In contrast, IdU tract lengths in fiber preparations from CPT-treated *ATM*^−/−^ cells were equivalent to untreated controls (Figure 3b,c). Predicting that the replication-stress resistant DNA synthesis evident in *ATM*-deficient cells was likely to be supported by PRIMPOL-dependent repriming, we also examined fibers from *ATM^−/−^PRIMPOL^−/−^*cells. Indeed, CPT-induced replication slowing was fully restored in *ATM^−/−^PRIMPOL^−/−^* cells (Figure 3c). Our results therefore show that under conditions of replication stress, an ATM-dependent regulation of replication slowing - presumably via fork reversal - typically counteracts PRIMPOL-dependent repriming at sites of DNA lesions. This explains why *ATM*-deficient cells accumulate PARP-activating post-replicative ssDNA gaps, which drives their hypersensitivity to PARPi.

Given BRCA1-A’s elevated recruitment to S-phase foci (Figure 3d,e) and CPT-stalled forks in ATM-deficient cells^30^, we hypothesized whether BRCA1-A contributes to fork slowing defects. Again, CPT-treated *ATM^−/−^*cells showed no IdU tract shortening upon treatments with CPT, while IdU-labelled fibers prepared from CPT-treated *ATM^−/−^ RAP80^−/−^*cells were shorter in length as in wild type cells (Figure 3f). The rescue of replication fork slowing in *ATM*-deficient co-disrupted for *RAP80* implicates BRCA1-A as a remodeler of lesion-stalled replication forks. By counteracting fork reversal, we surmise that BRCA1-A licenses PRIMPOL-dependent repriming and nascent-strand gap generation.

### Base excision repair intermediates cause replication defects in *ATM*-deficient cells

Since *ATM*-deficient cells accumulated nascent-strand gaps even under unchallenged conditions, we next questioned which DNA lesions might be triggering PRIMPOL-dependent replication repriming. Oxidative stress represents a primary driver of SSB formation in cells ^14^, and stimulates cell signaling and tolerance activities that are known to involve ATM ^28,43,44^. We thus performed experiments in which antioxidants were administered to probe whether oxidative stress influences steady-state DNA damage levels in *ATM^−/−^* cells. Indeed, β-mercapthoethanol (bME) or N-acetyl-cysteine (NAC) supplementation to cell culture media strongly attenuated spontaneous SSB levels in *ATM^−/−^* cells (Figure 4a). Similarly, these SSBs were suppressed in when cell culture oxygen concentrations were reduced from ambient levels of 20%, to physiological levels of 3% (Figure 4b), under which conditions nascent-strand ssDNA gap formation was also suppressed (Figure S5a). Physiological oxygen also dramatically increased the clonogenic survival of *ATM^−/−^*cells treated with PARPi (Figure 4c), with the enhanced PARPi resistance in *ATM-*knockout HCT-116 and RPE-1 cell lines manifesting as ∼17-20-fold increases in their IC_50_(olaparib) in cell viability assays (Figure 4d and Figure S5b). Furthermore, concordant ∼22-fold increases in the IC_50_(olaparib) were confirmed in ATMi treated cells (Figure S5c). Remarkably, this contrasted with the survival of HR-deficient cells (Figure 4c), where *BARD1*^Δ*/*Δ^ cells yielded equivalent dose-responses for olaparib in cell viability assays performed at 3% and 20% oxygen concentrations (Figure 4e). These findings confirm the specificity of oxygen-dependent replication defects in ATM-deficient cells.

**Figure 4:**
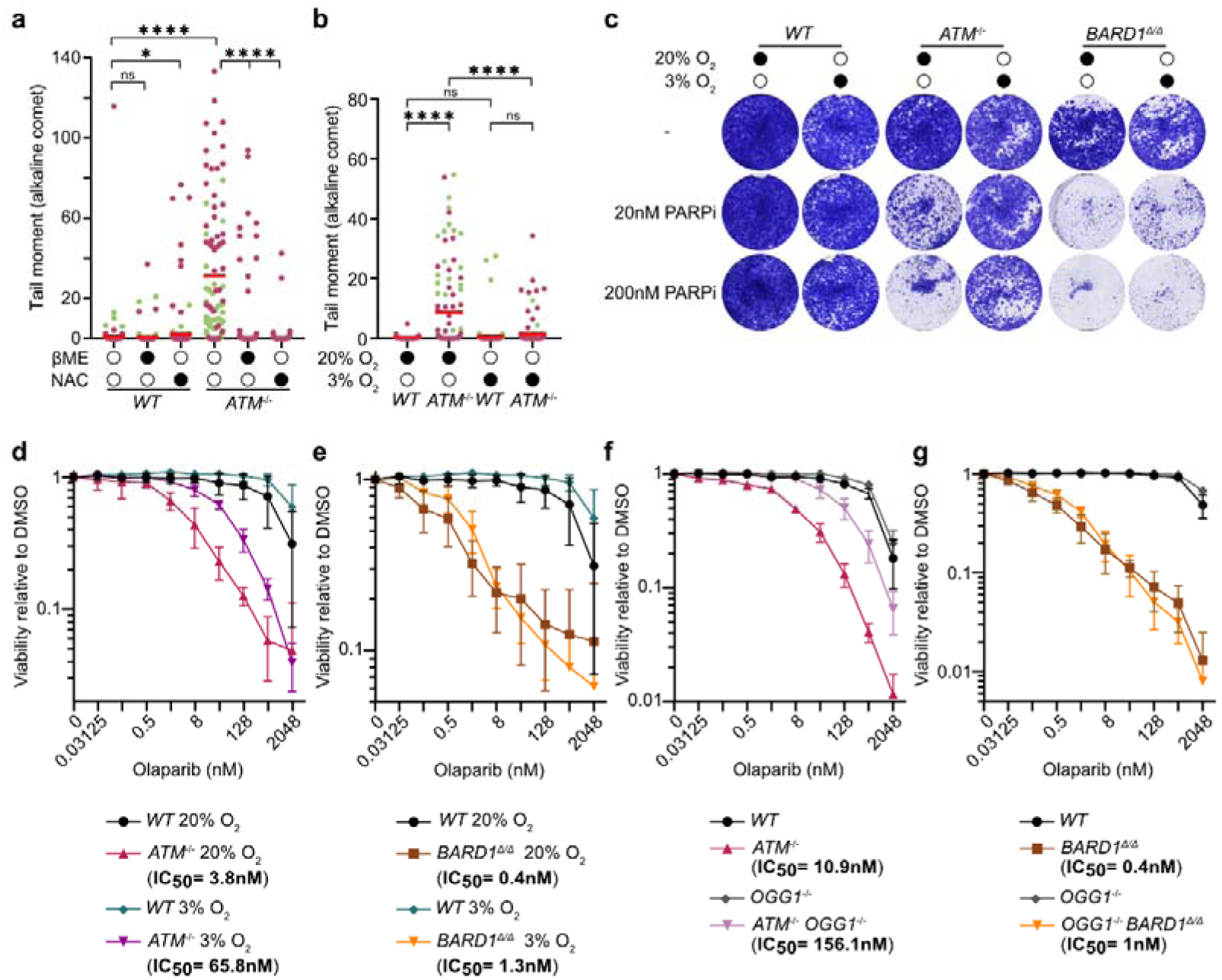
Base excision intermediates drive PARPi sensitivity in *ATM*-deficient cells. **a,b,** Alkaline comet assays of indicated HCT-116 cell lines cultured with 50 µM β-mercaptoethanol or 250 µM N-acetyl cysteine (NAC) for 72-96 h (**a**). Cells were cultured at ambient (20%) or physiological (3%) O_2_ conditions. Scatter plot data (**a**), *n=2* biological experiments (80–100 cells per experiment; each color denotes one replicate). Significance, two sided Kruskal-Wallis H test with Dunn’s correction for multiple comparisons. ****P ≤ 0.0001; ns, non-significant. **c,** Seven-day clonogenic survival of indicated *BARD1^AID/AID^* HCT-116 cell lines grown in the presence of indicated doses of olaparib (PARPi) under ambient (20%) or physiological (3%) O_2_ conditions. **d-g,** Resazurin survival assay of the indicated *BARD1^AID/AID^* HCT-116 cell lines grown for 7 days in the presence indicated doses of olaparib under ambient (20%) or physiological (3%) O_2_ conditions, *n=3* biological experiments, mean ± s.d. In experiments involving *BARD1*^Δ*/*Δ^ cells, all the indicated *BARD1^AID/AID^* cell lines were cultured in doxycycline (2 μg/ml) for 24 h before auxin (IAA, 1 mM) or dimethyl sulfoxide (DMSO) (carrier control), prior to olaparib addition. Throughout figure, *WT* denotes non-auxin-treated *BARD1^AID/AID^* cells.

The suppression of SSBs and PARPi toxicity in cultured *ATM^−/−^*cells upon oxygen manipulation led us to predict that polymerase stalling, either at a templated oxidized base adduct, or at an abasic (apurinic/apyrimidinic; AP) site generated upon adduct processing by a base excision repair (BER) glycosylase, could lead to post-lesion repriming by PRIMPOL. To explore these two scenarios, we considered the role of 8-Oxoguanine glycosylase 1 (OGG1), since 8-oxo-2′-deoxyguanosine (8-oxoG) is the major oxidized base adduct in cells ^45^. We therefore compared PARPi responses in *ATM^−/−^* and wild-type HCT-116 cell lines, and derivative clones in which both *OGG1* alleles were inactivated by CRISPR/Cas9-mediated mutagenesis. Strikingly, PARPi sensitivity was reduced dramatically in *OGG1^−/−^ ATM^−/−^* cells, when compared to its parental *ATM^−/−^* cell line (Figure 4f), with equivalent reductions in PARPi sensitivity also evident in *OGG1^−/−^* cells cultured with ATMi (Figure S5d). However, PARPi resistance was not seen in an *UNG^−/−^* HCT-116 cell line deleted for the Uracil-DNA glycosylase (UNG) BER glycosylase (Figure S5e), consistent with a specific exacerbation of the PARPi responses in *ATM*-deficient cells by endogenous 8-oxoG. By contrast, *OGG1*-deletion did not alter PARPi hypersensitivity in *BARD1*^Δ*/*Δ^ cells (Figure 4g). Considered together, this data confirms that while 8-oxoG derived AP sites make no discernable contribution to PARPi-induced toxicity in HR-deficient cells, they represent an important endogenous driver of replication-associated genomic instability and PARPi-induced synthetic lethality in *ATM-*deficient cells.

### SSBR sustains the viability of ATM-deficient cells

Cleavage of unrepaired AP sites in post-replicative ssDNA gaps can lead to chromosomal breakage during DNA replication, challenging genomic stability^46,47^. We therefore postulated that their frequent generation in *ATM^−/−^* cells would come at the expense of an acute reliance on post-replicative gap resolution activities. Given the hypersensitivity of ATM-deficient cells to PARPi, we first considered the role of the SSB repair pathway which is dependent on catalytic PARP1/2 activity^14^. Indeed, chemical inhibition of ATM in *PARP1^−/−^*RPE-1 cells triggered pronounced genomic instability, as evidenced by the induction of DSBs detected in neutral comet assays (Figure 5a), and consequent decreases in clonogenic cell survival (Figure 5b,c). Consistent evidence was additionally obtained using HCT-116 cells, where deletion of either PARP1, or its effector protein XRCC1, rendered cells equally susceptible to acute ATMi-induced toxicity (Figure 5d-f).

**Figure 5:**
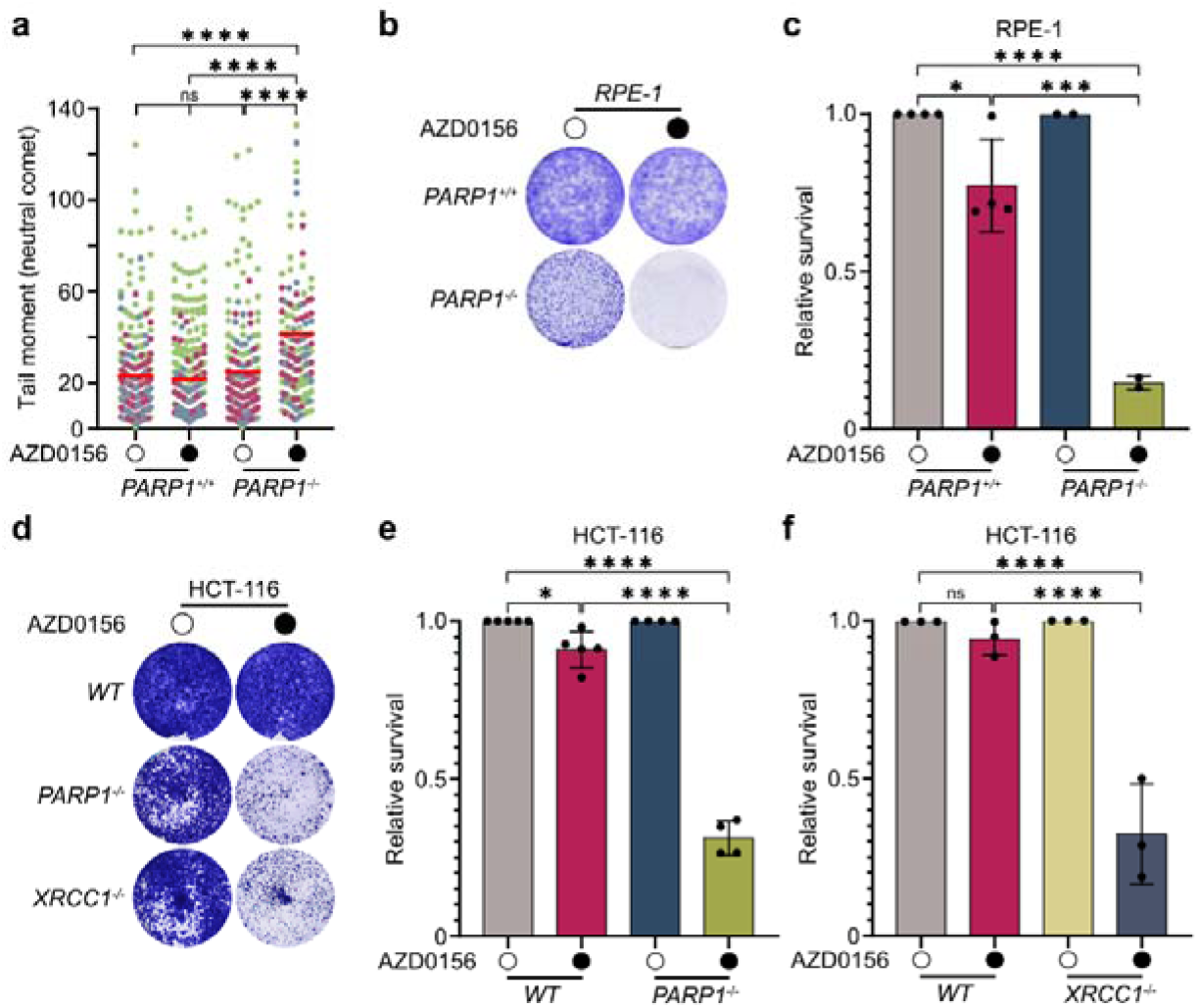
*ATM*-deficient cells are addicted to single-strand break repair. **a,** Neutral comet assay detection of DSBs with the indicated RPE-1 cell lines treated with AZD0156 (100 nM) or carrier (DMSO) for 48 h. Scatter plots, comet tail moments from *n=3* biological experiments (80-100 cells per experiment, each color denotes one replicate). Significance, two sided Kruskal-Wallis H test with Dunn’s correction for multiple comparisons. ****P ≤ 0.0001; ns, non-significant. **b,c,** Seven-day clonogenic survival assays for indicated RPE-1 cells treated with ATMi (AZD0156; 100 nM) or carrier (DMSO). Representative images **(b)** and quantification **(c)** of crystal violet staining. Data, mean ± s.d. of *n=2-4* biological experiments (each with >2 technical repeats). Significance, one-way ANOVA. ****P ≤ 0.0001; ***P ≤ 0.001; *P ≤ 0.05. **d-f,** Seven-day clonogenic survival assays with the indicated *BARD1^AID/AID^* HCT-116 cell lines supplemented with AZD0156 (ATMi; 500 nM) or carrier (DMSO, control). Representative images **(d)** and quantification **(e,f)** of crystal violet staining. Data, mean ± s.d. of *n=3-4* biological experiments (each with >2 technical repeats). Significance was determined by ordinary one-way ANOVA. ****P ≤ 0.0001; *P ≤ 0.05; ns, non-significant. Throughout figure, *WT* refers to non-auxin-treated *BARD1^AID/AID^* cells.

### HR mediates post-replicative nascent strand gap repair in ATM-deficient cells

In confirming the importance of SSB repair proteins in suppressing lethal genomic instability in *ATM*-deficient cells, we next considered the role of HR in repairing the frequent post-replicative ssDNA gaps accumulating in *ATM*-deficient cells (Figure 6a). DNA fiber analysis with staggered S1 nuclease treatments (30 min to 8 h post-IdU pulse) revealed that post-replicative ssDNA gaps were fully repaired within two hours in HR-proficient *ATM*^D*/*D^ *BARD1^AID/AID^* HCT-116 cells (Figure 6b). By contrast, when HR was inactivated with auxin during a 4-hour chase following the IdU pulse label, IdU fiber tracts remained fully-sensitive to S1 nuclease, signifying a complete loss of post-replicative gap repair (Figure 6c). To fully substantiate the role of HR in repairing nascent ssDNA gaps in *ATM*-deficient cells, we engineered an *BRCA2^AID/AID^* HCT-116 cell-line (and *ATM-*knockout derivatives), in which the sequences encoding mini-AID tag were introduced into the 3′ coding sequences of both copies of the *BRCA2* gene. Validating *BRCA2^AID/AID^* cells, we found that auxin treatments triggered rapid and efficient BRCA2 protein depletion (Figure S6a), inactivating HR as evidenced by the ablation of RAD51 recruitment and triggering acute hypersensitivity to olaparib treatment (Figure S6b,c). Monitoring the repair of S1 nuclease labile IdU fibers in *ATM^−/−^ BRCA2^AID/AID^* cells, our experiments again confirmed that nascent-strand gap repair was blocked by auxin-induced HR-inactivation (Figure 6c and Figure S6d).

**Figure 6:**
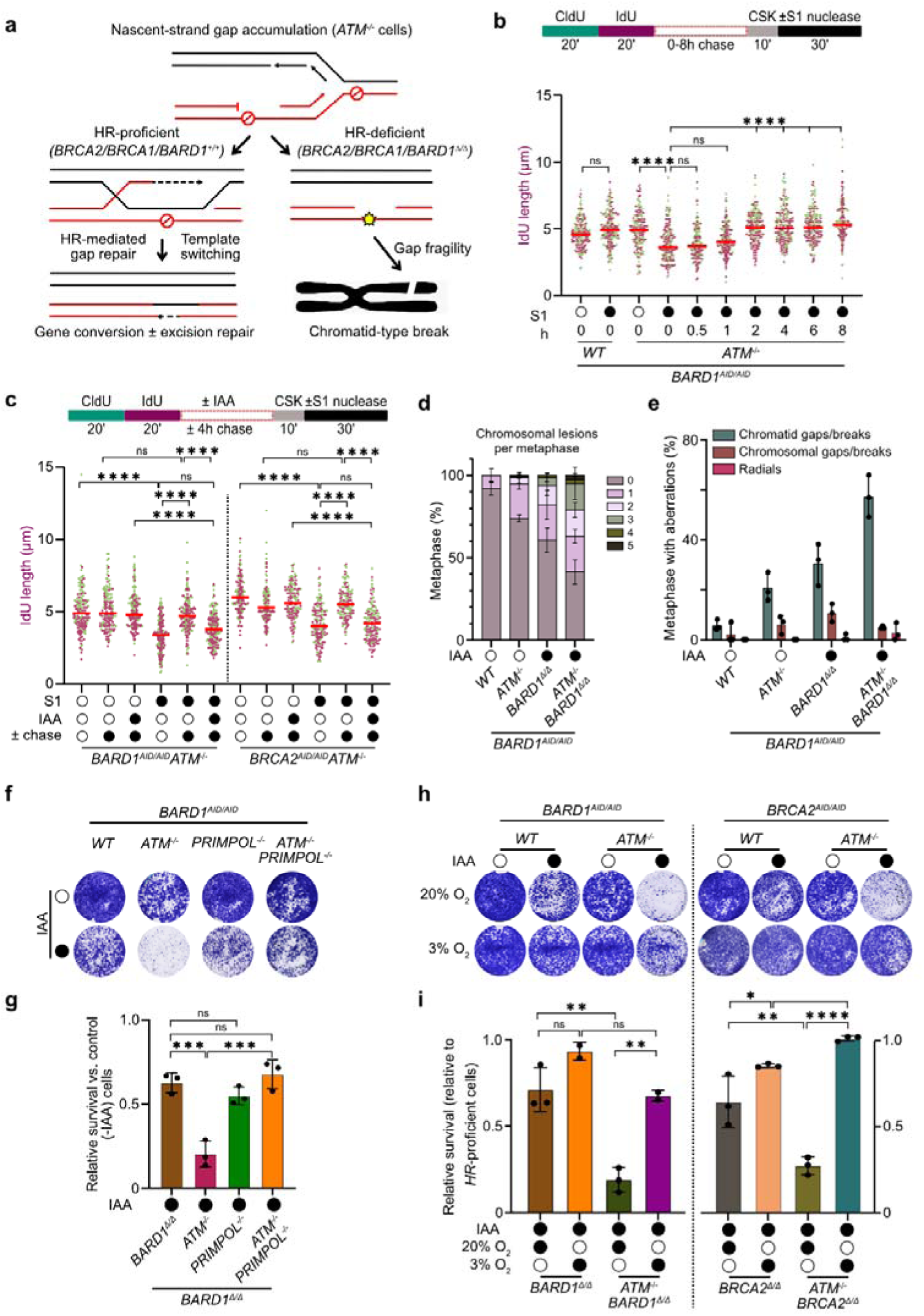
Cytotoxic nascent strand gaps drive HR addiction in *ATM*-deficient cells. **a,** Hypothesized role of HR-dependent post-replicative repair in *ATM*-deficient cells, and the predicted consequences of HR inactivation. **b,** Post-replicative gap repair kinetics in *ATM-*deficient cells. Schematic (top panel) depicting nascent-strand gap detection by S1-nuclease coupled DNA fiber assay. Following the IdU pulse, a 0-8 h chase was performed with thymidine-supplemented medium prior to S1nuclease treatment. Scatter plots (bottom panel) showing IdU tract lengths in DNA fibers. Horizontal bars, median values of >250 fibers per cell line in *n=2* biological experiments, each represented in a different color. Significance was determined by two sided Kruskal-Wallis H test with Dunn’s correction for multiple comparisons. ****P ≤ 0.0001; ns, non-significant. **c,** HR repairs nascent strand gaps in *ATM-*deficient cells. Schematic (top panel) depicting nascent-strand gap detection by S1-nuclease coupled DNA fiber assay. As in **b**, but with the addition of auxin or DMSO (mock) during the 4 h chase. Scatter plots (bottom panel) showing IdU tract lengths in DNA fibers from the indicated *BARD1^AID/AID^* or *BRCA2^AID/AID^*HCT-116 cells lines. Horizontal bars, median values of >250 fibers per cell line in *n=2* biological experiments, each represented in a different color. Significance was determined by two sided Kruskal-Wallis H test with Dunn’s correction for multiple comparisons. ****P ≤ 0.0001; ns, non-significant. **d,e,** Chromosome lesions in metaphase chromosome prepared from the indicated *BARD1^AID/AID^* cell lines. Quantification (**d**) and classification (**e**) of chromosome aberrations. Data, *n=3* biological experiments, each comprising >40 metaphases. Significance, ordinary one-way ANOVA test. ****P□<□0.0001; ***P□<□0.0005; **P□<□0.005; *P□<□0.05. **f-i**, Seven-day clonogenic survival assays with the indicated *BARD1^AID/AID^* and *BRCA2^AID/AID^* cell lines (**f,i**). Cells grown at ambient (20%) or physiological (3%) O_2_ conditions. Representative images (**f,h**) and quantification (**g,i**) of crystal violet staining. Data, mean ± s.d. of n=3 biological experiments (each with >2 technical repeats). Significance was determined by ordinary one-way ANOVA with Tukey’s correction for multiple comparisons. ****P ≤ 0.0001; ***P ≤ 0.001; **P ≤ 0.01; * P ≤ 0.05; ns, non-significant. Throughout the figure, *WT* refers to non-auxin-treated parental *BARD1^AID/AID^* or *BRCA2^AID/AID^* cells. In the experiments involving *BARD1*^Δ*/*Δ^ and *BRCA2*^Δ*/*Δ^ cells, all the indicated *BARD1^AID/AID^* and *BRCA2^AID/AID^* cell lines were cultured in doxycycline (2 μg/ml) for 24 h before auxin (IAA, 1 mM) or dimethyl sulfoxide (DMSO) (carrier control).

### Post-replicative gap repair renders *ATM*-deficient cells addicted to HR

In confirming HR as the major mechanism repairing nascent strand gaps in *ATM*-deficient cells (Figure 6a), we next investigated the consequences of chronic inactivation of this repair in *ATM-*knockout cells and their controls. As before (Figure 1g), neutral comet assays only detected spontaneous DSBs in HR-deficient (auxin-treated) *BARD1*^Δ*/*Δ^ cells but not in *ATM*^−/−^ cells (Figure S7a). However, HR-inactivation in *ATM^−/−^ BARD1*^Δ*/*Δ^ cells led to a strong induction of DSBs, with signals surpassing that seen in parental *BARD1*^Δ*/*Δ^ cells (Figure S7a). The synergistic effect of inactivating HR in *ATM*-deficient cells on genome stability was also evident upon quantifying chromosomal lesions in metaphase chromosome spreads. Around 60% of metaphases prepared from *ATM^−/−^ BARD1*^Δ*/*Δ^ cells harbored between 1 and 5 chromosome lesions (Figure 6d and Figure S7b), with these mainly consisting of chromatid-type breaks (Figure 6e), the predicted product of a post-replicative breakage or cleavage event at an unrepairable nascent-strand gap (Figure 6a). Profound chromosomal instability in *ATM^−/−^ BARD1*^Δ*/*Δ^ cells was already apparent 48h after auxin addition and coincided with an induction of apoptosis in up to 40% cells by 96h (Figure S7c,d). Finally, clonogenic cell survival assays confirmed synthetic lethality in cells co-inactivated of BARD1 and ATM (Figure S7e). In these experiments, cell viability could be completely restored upon re-expression of wild type BARD1, but not an HR-deficient mutant of BARD1^24^ (BARD1^D712A^; Figure S8a,b), reconfirming the importance of HR for the viability of *ATM*-deficient cells^22,48^.

Given the extreme toxicity inflicted on *ATM*-deficient cells upon BARD1 inactivation, we hypothesized that this likely stemmed from an inability to repair nascent-strand gaps after DNA replication using an HR-dependent template switching mechanism (Figure 6a). We reasoned that if this was correct, the acute reliance of *ATM*-deficient cells on HR would be relieved by either blocking nascent-strand gap formation, or by reducing the oxidative base damage priming their generation during replication. To block gap formation, we tested the impact of PRIMPOL-deletion in *ATM^−/−^ BARD1^AID/AID^* cells on cell survival following auxin-induced inactivation of HR. In these experiments, auxin-treatments reduced clonogenic survival in both *ATM^−/−^ BARD1^AID/AID^* and *PRIMPOL^−/−^ BARD1^AID/AID^* cell lines, indicative of acute HR-reliance characterizing both *ATM* and *PRIMPOL*-deleted cells (Figure 6f,g). However, auxin-induced toxicity, and associated cell proliferation defects (Figure 6f,g), were largely alleviated in *ATM^−/−^ PRIMPOL^−/−^ BARD1^AID/AID^* cells, aligning with HR’s importance in repairing post-replicative ssDNA gaps in *ATM*-deficient cells. To manipulate lesion abundance, we repeated clonogenic assays that compared the survival of *ATM^−/−^* cells co-inactivated for HR at physiological and ambient oxygen concentrations. Here, physiological oxygen strongly suppressed the synthetic lethal effect of inactivating HR in both *ATM^−/−^ BARD1*^Δ*/*Δ^ and *ATM^−/−^ BRCA2*^Δ*/*Δ^ HCT-116 cell-lines (Figure 6h,i), and similarly in *BARD1*^Δ*/*Δ^ and *BRCA2*^Δ*/*Δ^ cells treated with ATMi (Figure S8c,d). Finally, clonogenic survival assays demonstrated a concordant dramatic attenuation of ATMi-induced toxicity in *BRCA1^−/−^ TP53^−/−^* RPE-1 cells when experiments were performed at physiological oxygen (Figure S8e,f). Considered together, our data reveals that spontaneous oxidative base adducts drive pathological accumulation of nascent strand gaps in replicating *ATM*-deficient cells. These lesions also provide an explanation for their hypersensitivity to PARPi, and a rationale for their excessive demands on post-replicative repair by HR.

## Discussion

Altogether, our findings define the mechanism underpinning PARPi-induced killing of ATM-deficient cells, revealing a pivotal role for ATM in regulating replication dynamics at endogenous base lesions. This work resolves a long-standing synthetic lethal interaction first reported nearly two decades ago^4,5^. Unexpectedly, the mechanism diverges sharply from the model of PARPi-induced synthetic lethality in *BRCA*-deficient tumors, where replication fork collisions with trapped PARP1/2 complexes generate DNA damage that overwhelms HR-deficient cells^3^. Instead, in ATM-deficient cells, PARPi toxicity arises from discontinuous leading-strand synthesis during replication, a pathological outcome driven by unrestrained PRIMPOL-dependent repriming at endogenous base adducts that stall DNA synthesis.

To understand the cause of stochastic repriming in ATM-deficient cells, we show that ATM activity enforces replication fork slowing, most likely through fork reversal, at sites of replication-blocking base adducts. Loss of ATM disrupts this control, driving excessive PRIMPOL-dependent bypass. The mechanism by which ATM orchestrates replication slowing and reversal remains incompletely defined, but our data implicate ATM-dependent inhibition of BRCA1-A complex recruitment to lesion-stalled forks. Consistent with this, the BRCA1-A complex both prevents fork slowing in ATM-deficient cells exposed to CPT-induced replication stress and sustains replication repriming at spontaneous oxidized base adducts. We propose that ATM-mediated phosphorylation events restrict BRCA1-A accumulation at stalled forks, thereby coupling DNA damage signalling with replication control. This model is supported by proteomic analyses of nascent chromatin showing that ATM catalytic activity limits BRCA1-A at human replisomes under chronic replication stress^30^. It also explains prior observations that BRCA1-A contributes to PARPi and CPT toxicity in ATM-deficient contexts^32^ and compromises proliferative fitness upon catalytic ATM inhibition^33^.

We suggest that ATM-mediated fork slowing evolved to prevent transmission of unrepaired base adducts on the leading-strand template to daughter chromatids. The persistence of such lesions - exposed within unreplicated single-stranded DNA - poses a high risk of chromatid breakage and acentric fragment formation, events that promote genomic instability. The severe chromatid breakage and cell death that occurs in ATM-deficient cells when post-replicative repair by either SSBR or HR was inactivated is consistent with this model.

Our discovery that endogenous base lesions - particularly intermediates of 8-oxoG excision repair - drive replication dysfunction in ATM-deficient cells may hold relevance to A-T pathophysiology. In *Atm*-knockout mice, endogenous reactive oxygen species (ROS) progressively erode hematopoietic stem cell (HSC) functionality, leading to bone marrow failure^49^. Impaired replication-coupled resolution of oxidized base repair intermediates could potentially contribute to this attrition of HSCs in A-T. In post-mitotic neurons undergoing active DNA demethylation, TET enzyme–mediated hyper-oxidation of methyl-CpG dinucleotides at neural enhancers stimulates recurrent cycles of BER^50^. It is conceivable that similar processes in ATM-deficient neural stem and progenitor cells could generate oxidative DNA intermediates that exacerbate replication stress, thereby contributing to neurodevelopmental decline. Although oxidative stress has long been implicated in A-T pathogenesis, the extent to which pathology reflects redox imbalance versus defective DNA damage signaling remains unresolved^17,51^. By establishing a direct mechanistic connection between oxygen-induced DNA damage and ATM-dependent replication control, our findings provide a potential unifying explanation for these observations.

## Methods

### Cell Culture

The genetically engineered *BARD1^AID/AID^* HCT-116 colorectal cancer cell-line, that also harbors bi-allelic targeted integrations of a doxycycline-inducible *OsTIR1* transgene at the *AAVS1* locus (using Addgene plasmid 72835 ^52^), was generated and described previously ^24,53^. *BRCA2^AID/AID^* HCT-116 cells were prepared from the same parental *AAVS1^iOsTIR^*^1^*^/iOsTIR^*^1^ HCT-116 cell-line. Briefly, donor plasmid for endogenous AID tagging of BRCA2 C-terminus was prepared by overlapping fusion PCR of 3 fragments: 1) a left homology arm amplified from HCT-116 genomic DNA using oligos BRCA2_LHA_F: (CAATAGCTGACGAAGAACTTGC) and BRCA2_LHA_R: (TCTTTAGGACAAGCACTCTTCTCCTTGGCGCCTGCACCGGAACCTGAACCGATATATTTTTTAGTTGTAATTGTGTCC), 2) a right homology arm amplified from HCT-116 genomic DNA using oligos BRCA2_RHA_F: (AAGTTATTAGGTCCCTCGAAGAGGTTCACTAGGATCCGGTACCCACAAATGCGACAATAAATTATTGAC) and BRCA2_RHA_R: (CCGAGCTCAGCCAAAGATG), and 3) an AID tag and hygromycin selection cassette amplified from pMK287 (Addgene plasmid #72827) (PMID 27052166) using oligos mAIDcassette_F: (GGTTCAGGTTCCGGTGCAGGCGccAAGGAGAAGAGTG) and mAIDcassette_R: TGGGTACCGGATCCTAG. Fused product was ligated into pJET 1.2 by blunt end cloning following the CloneJET PCR Cloning Kit manufacturer’s protocol.

All knockout/knockin cell lines were prepared by CRISPR–Cas9. The following gene-specific gRNAs were cloned into BbsI digested-pSpCas9(BB)-2A-GFP (pX458) (48138, Addgene): *ATM* (TGATAGAGCTACAGAACGAA); *PRIMPOL* (GTTGCAGTAGAAACCATTGA, CAATCTTGGTCTATACACTG); *PARP1* (AAGAAGACAGCGGAAGCTGG); *XRCC1* (AGACACTTACCGAAAATGGC), *OGG1* (TGGCTCAACTGTATCACCAC, ACACTGGAGTGGTGTACTAG), *UNG* (CAAAGCCCACGGGCACGTTG), *BRCA2* (ATATATCTAAGCATTTGCAA), *ABRAXAS* (TGATCTGATCTGAATGACGA for KO and AGGGTTTTGGTGAATATTCA for KI) and *BRCC36* (CGCTCTGAGCACAGAGAAGG for KO and GCCAAACAGTTATATGAGGA for KI). 2 μg of corresponding pX458 plasmid was electroporated into 1×10^6^ cells using a Lonza 4D-Nucleofector according to the manufacturer’s protocol designed for HCT-116 cells (SE buffer, program EN-113). To prepare knockin *ABRAXAS^S^*^406^*^A/S^*^406^*^A^* and *BRCC36^QSQ/QSQ^* cells were co-transfected with homology directed repair template (ssODN). Cell-sorts were performed using a Sony MA900 sorter, into medium containing 40% FBS and recovered for 4 days post-sort. Cells were then seeded at low density (250, 500, 1000 cells per 10 cm dish). After 7-10 days, individual clones were picked and expanded in 24-well plates. Successfully edited clones were identified by native PAGE analysis, by comparing CRISPR/Cas9 target site-spanning short PCR amplicons amplified from genomic DNA prepared from targeted and parental (control) cell clones. Gene knockouts were then validated by immunoblotting and Sanger sequence verification of gene-disruptive indels. The mycoplasma-free status of all cell lines was first verified upon entering the laboratory, and then subsequently reverified in routine tests.

To generate lentivirus for stable transgene complementation (RAP80^WT^ and RAP80^UIM1/2^), HEK293T female embryonic kidney cells (obtained from Francis Crick Institute Cell Services; RRID: CVCL_0063) were co-transfected with a lentiviral vector encoding the transgene of interest, pHDM-tat1b, pHDM-G, pRC/CMV-rev1b and pHDM-Hgpm2 using 1.29 μg polyethylenimine per μg of DNA in Opti-MEM (31985062, Thermo Fisher). Viral supernatants were collected at 48 h and 72 h after transfection, syringe-filtered (0.45 μm), and immediately used to transduce target cells populations in the presence of 4 μg/ml polybrene. Transduced populations were then prepared by neomycin selection.

In experiments that either included, or contrasted, cellular responses to *BARD1*^Δ*/*Δ^ (BRCA1/BARD1-depleted) cells, all cells were treated identically, plating in media with 2 µg/ml doxycycline for 16-24 h prior to either treatment with auxin (IAA, 1 mM), or mock (DMSO), for at least 1 h before experiments began. Parental cell-lines, including described HCT-116, RPE-1, MDA-MB-231 cell-lines, and genetically altered derivative cell-lines were maintained in Dulbecco’s modified Eagle medium (DMEM)–high glucose (41966052, Gibco) supplemented with 10% FBS, 1% penicillin–streptomycin and 2 mM L-glutamine. Cells were cultivated at 37°C with 5% CO_2_ and 20% O_2_, for some of the experiment, cells were grown in 3% O_2_ condition.

*RAP80^−/−^ BARD1^AID/AID^* and *BARD1^AID/AID^* HCT-116 complemented with control (GST) or BARD1 transgenes (wild type and BARD1^D712A^) were prepared and described previously ^24^. The RPE-1 *FEN1*^−/−^ cells ^25^ were provided by K. Caldecott, RPE-1 *PARP1*^−/−^ cells were provided by N. Lakin, RPE-1 *ATM*^−/−^ cells were provided by F. Cortes-Ledesma, and *TP53^−/−^BRCA1^−/−^* RPE-1 FRT cells were provided by S.P. Jackson ^54^.

### Mice

*Atm^−/−^* mice (allele ID *Atm^tm1Awb^*; MGI ID 1857132) bred on a C57BL/6 background were generated and described elsewhere ^55^. The production and breeding of genetically altered mice was carried out in accordance with the UK Home Office Animal (Scientific Procedures) Act 1986, with procedures reviewed by the Clinical Medicine Animal Welfare and Ethical Review Body at the University of Oxford and conducted under project license PP8064604. Animals were housed in individually ventilated cages, provided with food and water ad libitum and maintained on a 12-h light–dark cycle (150–200□lx). The only reported positives on Federation of European Laboratory Animal Science Associations health screening over the entire time course of these studies were for Helicobacter hepaticus and Entamoeba spp. Experimental groups were determined by genotype and were, therefore, not randomized, with no animals excluded from the analysis. Sample sizes for fertility studies were selected on the basis of previously published studies and all phenotypic characterization was performed blind to the experimental group. All mice used in this study were generated on or backcrossed onto a C57BL/6 background (>5 generations).

The experiments involved age-matched 8–16-week-old male or female animals on an inbred C57BL/6 background. Mice of a certain genotype were selected using a unique mouse identifier that does not indicate mouse genotype; thus, phenotype–genotype relationships were determined only at the data analysis stage. All experiments were approved by the University of Oxford Ethical Review Committee and performed under a UK Home Office License in compliance with animal use guidelines and ethical regulations.

### Survival and proliferation experiments

#### Resazurin viability assay

To measure cell viability *BARD1^AID/AID^* HCT-116 cells were plated in the corresponding cell numbers (*BARD1^AID/AID^* (untreated with auxin), *RAP80^−/−^, ABRAXAS^−/−^, BRCC36^−/−^,PRIMPOL^−/−^, UNG^−/−^* and *OGG1^−/−^:* 300, *BARD1*^Δ*/*Δ^: 500, ATMi/*ATM^−/−^* and *ATM^−/−^RAP80^−/−^, ATM^−/−^ABRAXAS^−/−^, ATM^−/−^BRCC36^−/−^, ATM^−/−^ OGG1^−/−^:* 500, and *ATM^−/−^ PRIMPOL^−/−^:* 1000) and in the media containing 2 µg/ml doxycycline in triplicate in a 96-well plate. After 24 h, 1 mM auxin (IAA, 3-indoleacetic acid; I2886, Merck) or corresponding amount of DMSO was added. One hour after IAA addition, olaparib (S1060, Selleckchem) or cisplatin (P4394, Merck) was added to the indicated final concentrations. 1×10^2^ RPE-1 cells were seeded per well in 96-well plate in three technical repeats, incubated overnight and then cells were treated with indicated concentration of olaparib. Seven days after drug addition, the medium was replaced with phenol red-free DMEM (21063-029, Gibco) supplemented with 10% FBS, penicillin–streptomycin, 2 mM l-glutamine and 10 μg/ml resazurin (R7017, Merck). Following 3-4 hours incubation and resazurin color modification, relative fluorescence was measured at 595 nm with a CLARIOstar plate reader (BMG Labtech). The mean of three technical repeats after background subtraction was taken as the value for a biological repeat and the survival curves are plotted as the mean of three biological experiments ± s.d.

#### Survival analysis by crystal violet staining

1×10^4^ cells were seeded per well of a 12-well plate in technical duplicates for each cell line in the presence of doxycycline (2 μg/ml). After 24 h, IAA (1 mM) or carrier (DMSO) was added. 5×10^3^ RPE-1 cells were seeded per well of 6-well plate. In the experiment to measure PARPi sensitivity, olaparib was added to the indicated final concentrations 1 h after IAA treatment. After 7 days of growth, medium was replaced by crystal violet staining solution (0.5% crystal violet in 25% methanol). Cells were stained for 10-15 min, washed several times with ddH_2_O and left to dry before scanning. To quantify the relative viability, the stain was dissolved in 2% SDS, diluted 1:4 with H_2_O and the absorbance was measured at CLARIOstar plate reader (BMG Labtech) at 575 nm. Representative wells were selected for display.

#### Growth/proliferation assay

2×10^5^ cells were seeded per well of a 6-well plate in the presence of doxycycline (2 μg/ml). After 24 h, IAA (1 mM) or carrier (DMSO) was added. Cells were grown for 3 days, then counted and 2×10^5^ cells were re-plated. The process was repeated over 9-day period.

### Immunofluorescence

Cells were exposed to 2 µg/ml of doxycycline and 24 hours later, 2×10^5^ cells were run through a 70-μm mesh cell strainer and seeded on 2 fibronectin-coated glass coverslips (13 mm) in a single well of a 6-well plate. The growth medium contained doxycycline (2 μg/ml) and either DMSO or 1 mM final concentration of auxin (IAA). For the experiment analyzing RAD51 foci, 24 h following IAA addition, cells were either treated with 500 nM olaparib for 24 h or irradiated (5 Gy) + 2 h recovery and fixed in 2% PFA. In experiments analyzing nuclear PAR levels, cells were treated with 10 µM PARGi inhibitor (SML1781, Merck) for total 45 min before fixation with 4% PFA. Coverslips with fixed cells were first blocked (3% BSA, 0.1% Triton X-100 in PBS) for 15 min at room temperature (RT), followed by 1 h (2 h for PAR staining, overnight incubation at 4□C for RAP80 staining) incubations with primary antibody in a humidity chamber. In experiments analyzing replication or transcription, cells were treated with 10 µM 5-ethynyl-2’-deoxyuridine (EdU, ab146186, Abcam) or 200 µM 5-ethenyl uridine (EU, E10345, Life Technologies) prior to fixation. Click-iT EdU Cell Proliferation Kit, Alexa Fluor 647 (C10340, Life Technologies) was used to label EdU/EU-positive cells according to the manufacturer’s protocol between blocking and primary antibody incubation steps. The following primary antibodies were used: rabbit anti-RAD51 (1:1,000, 70-001, BioAcademia), mouse anti-γH2AX (1:1,000, 05-636, Millipore), rabbit anti-RAP80 (1:600, A300-763A, Bethyl) and mouse anti-PAR (1:500, 4335-MC-100, BioTechne). Next, coverslips were washed 3 times with PBS + 0.1% Triton X-100 before incubation with secondary antibodies for 1 h at room temperature in a humidity chamber. The following secondary antibodies were used: goat anti-mouse Alexa Fluor 488 (1:500, A-11001, Invitrogen), and goat anti-rabbit Alexa Fluor 568 (1:500, A-11011, Invitrogen). Coverslips were then washed 3 more times with PBS + 0.1% Triton X-100 and mounted on glass microscope slides using a drop of ProLong Gold antifade reagent with DAPI (P36935, Life Technologies). Images were acquired on a Leica DMi8 wide-field microscope. CellProfiler (Broad Institute) was used for foci quantification. Images were visualized and saved in Fiji and assembled into figures in Affinity Designer.

### DNA fiber assays

DNA fiber assays were performed essentially as previously described ^56^. Briefly, cells were pulse-labeled with 30 µM CldU and 250 µM IdU (C6891 and I71250, MP Biomedicals) in sequential 20 min incubations at 37°C. Cells were washed between and after the pulses with warm medium, and following the IdU pulse, cells were scraped into cold 0.1% BSA/PBS and centrifuged at 7000 rpm for 5 min at 4°C. The pellet was resuspended in cold 0.1% BSA/PBS, 2 µl of the cell suspension were spotted on glass slide and lysed with 7 µl of lysis solution (50 mM EDTA; 200 mM Tris-HCl pH 7.5, 0.5% SDS) for 3-5 minutes. The slides were tilted ∼45° relative to horizontal surface allowing the spreading of DNA fibers. Slides were air-dried overnight before fixation with 3:1 fixative methanol: acetic acid solution for 10 min at room temperature and incubation under denaturating conditions of 2.5 M HCl for 80 minutes. Slides were then washed with PBS and blocked with 5% BSA for 20 minutes at room temperature. DNA fibers were stained with the following antibodies: 2 h incubations with primary antibodies mouse anti-BrdU (1:25; 347580, BD Biosciences) and rat anti-BrdU (1:400; ab6326, Abcam); 1 h incubations with secondary antibodies rabbit anti-mouse (1:500; A11061, Invitrogen) and chicken anti-rat (1:500; A21470, Invitrogen); 1 h incubations with tertiary antibodies goat anti-rabbit (1:500; A11036, Invitrogen) and goat anti-chicken (1:500; A11039, Invitrogen). All immunostaining was performed at RT in a humidified chamber protected from light with the PBS washes between each immunostaining. Slides were then mounted in Prolong Gold antifade mounting medium (P36935, Life Technologies) and kept at 4°C prior imaging on a Leica DMi8 wide-field microscope. Both acquisition and analysis were performed blindly using ImageJ software and 75-100 fibers were analyzed per sample per experiment.

To monitor DNA damage-induced slowing of DNA synthesis, cells were treated with 100 nM camptothecin (CPT; 11694, Cayman) throughout the duration of IdU pulse.

In S1-nuclease coupled fiber assays, following the IdU pulse, cells were permeabilized with cytoskeletal buffer (10 mM PIPES pH 6.8; 100 mM NaCl; 300 mM sucrose; 1 mM EGTA; 1 mM MgCl_2_; 1 mM DTT supplemented with protease inhibitor) for 10 minutes at RT. CSK buffer was gently removed and cells were washed with PBS and 1X S1 buffer (30 mM sodium acetate pH 4.6, 1 mM zinc acetate, 5% glycerol). Then, one half of cells were treated with 20 U/ml of S1 nuclease diluted in 1X S1 buffer (18001016, Invitrogen), and the second half were kept only in 1X S1 buffer for 30 min at 37°C. The S1 buffer was removed and cells were scraped into cold 0.1% BSA/PBS and centrifuged at 7000 rpm for 5 min at 4°C and proceed as described above. In the experiment measuring gap repair, following IdU labelling cells were washed twice with media and allowed to recover in the complete media supplemented with 500µM thymidine and either auxin (IAA) or DMSO for 0, 0.5, 1, 2, 4, 6 and 8h. Samples were than prepared as above for S1 DNA fiber assay.

### Cytological and sister chromatid exchange assays (SCEs)

Metaphase spreads were prepared by standard protocol. Briefly, cells were exposed to 50 ng/ml KaryoMAX colcemid (15210040, Gibco) for 3 h prior collecting both detached and trypsin-treated cells. The cells were then centrifuged, resuspended in 75 mM KCl and incubated for 15□min at 37°C before being fixed in ice cold Carnoy’s fixative (3:1 methanol: glacial acetic solution). Approximately 20 μl of cell suspension was dropped onto clean slides and left to dry overnight. The slides were then stained and mounted with Prolong gold DAPI mounting medium (P36935, Life Technologies). The slides were acquired using Leica DM660 B microscope equipped with a Cytovision software (Leica).

To image and quantify sister chromatid exchanges (SCEs), slides were airdried for 2 days prior staining with 10 µg/ml Hoechst 33258 for 30 minutes in dark at RT. After washing in Sorenson Buffer (1:1 mix of 100 mM Na_2_HPO_4_ and 100 mM NaH_2_PO_4_), slides were exposed to UV light (354 nm) while heated to 55°C for 10 minutes, then incubated in 1X SSC (S6639, Sigma) buffer at 50°C for 1 h, and finally stained with 10% KaryoMAX Giemsa solution (10092013, Gibco). After washing in ddH_2_O and mounting in Mowiol mounting medium, images were acquired using Olympus BX60 microscope and cellSens software (Olympus). A minimum of 40 metaphases were processed and analyzed blindly per each cytogenic experiment with ImageJ software.

### Comet assays

Treated cells were trypsinized, counted and resuspended in ice-cold PBS at concentration 2×10^5^ cells/ml. Cell suspensions were mixed with equal amounts of melted 1.2% low melting agarose, layered on agarose-coated slides and kept in 4°C until the agarose plugs solidified. The cells were lysed with cold lysis buffer (2.5 M NaCl, 100 mM EDTA pH 8.0, 10 mM Tris-HCl pH 10, 1% Triton X100 and 1% DMSO) for 1 h (alkaline comet assay), or overnight (neutral comet assay). For alkaline comet assays, slides were equilibrated in cold running buffer (1 mM EDTA pH 8.0, 50 mM NaOH and 1% DMSO) for 45 minutes prior the electrophoresis performed for 15 min at 25 mA at 4°C. Slides were then neutralized in a 0.4 M Tris-HCl buffer pH 7.0 for 1 h. For the neutral comet assays, slides were equilibrated in 1X TBE and electrophoresis was performed at 25 min at 5 mA at 4°C. Slides were then dehydrated in 70% EtOH, dried at 37□C and stained with 25 µM propidium iodide before acquiring images on a Leica DMi8 wide-field microscope. At least 50 nuclei were acquired per sample per biological experiment. The analysis was performed blindly by measuring tail moment with the OpenComet plugin for Fiji software. Individual tail moments were plotted as scatter plot graphs in Prism 10 (GraphPad software).

### Western blotting

Cells were washed once with PBS and lysed by resuspension in ice-cold benzonase cell lysis buffer (40 mM NaCl, 25 mM Tris pH 8.0, 0.05% SDS, 2 mM MgCl_2_, 10 U/ml benzonase, and cOmplete EDTA-free protease inhibitor cocktail). Extracts were then incubated on ice for 10 min and mixed with Laemmli buffer, boiled at 95°C for 5 min before loading on SDS–PAGE gels. After transfer, membranes were blocked with 5% milk in PBST for at least 30 min and then incubated overnight with primary antibody in PBST supplemented with 0.03% NaN3 and 3% BSA. Primary antibodies used for western blot in this study include: rabbit anti-BRCA2 (1:1000, OP95, Merck), mouse anti-β-actin (1:2000, A1978, Sigma-Aldrich) and rabbit anti-OsTIR1 (a gift from M. T. Kanemaki; 1:1000). Following primary, membranes were incubated with either HRP-conjugated goat anti-mouse (1:20000, 62-6520, Thermo Fisher) or HRP-conjugated goat anti-rabbit (1:20000, 65-6120, Thermo Fisher) secondary antibodies. Membranes were developed with Clarity Western ECL Substrate (Bio-Rad, 170-5061) and imaged using a Gel Doc XR System (Bio-Rad).

### B cell isolation and culture

B cells were purified from red blood cell-lysed single-cell suspensions of mouse spleens by magnetic negative selection using a B cell isolation kit (130-090-862, Miltenyi Biotec). B cells (5×10^5^ per well in a 6-well plate) were cultured in RPMI supplemented with 10% fetal calf serum (FCS), 100□U/ml penicillin, 100□ng/ml streptomycin, 2□mM L-glutamine, 1x MEM nonessential amino acids, 1□mM sodium pyruvate and 50□μM β-mercaptoethanol. B cells were stimulated with 10□ng/ml mouse recombinant interleukin 4 (214-14-20, Peprotech) and agonist anti-CD40 antibody (0.5□μg/ml; FGK45.5 Miltenyl Biotec). Cultures were grown at 37°C with 5% CO_2_ under ambient oxygen conditions. B cells were harvested at both T=0 h and following 72 h stimulation with cytokines cocktails for comet assay.

### Apoptosis FACS staining

5×10^5^ cells were collected, washed with PBS and resuspended in 100□µl 1X Annexin binding buffer (10 mM HEPES pH 7.4, 140 mM NaCl, 2.5 mM CaCl_2_), before staining with FITC-Annexin V (1:50, 556420, BD Pharmingen) antibody and 1□mg/ml propidium iodide (PI, 1:100, P4170, Merck). After an incubation for 15□min at room temperature in the dark, topped up with 400□µl 1X Annexin binding buffer, cells were analyzed on the flow cytometer (Attune NxT flow cytometer, ThermoFisher) immediately. The data were analyzed with FlowJo software version 10 (Tree Star).

### Statistics

Prism 10 (Graphpad Software) was used for graph preparation and statistical analysis. Relevant statistical methods for individual experiments are detailed within figure legends.

## Supporting information

Complete Supplementary Figures 1-8

## Acknowledgments

We thank all members of the laboratory of J.R.C. for discussions and support; E. Gogola for critical feedback on the manuscript. F. Cortes Ledesma (Universidad de Sevilla), N. Lakin (University of Oxford), K. Caldecott (University of Sussex), S. Jackson (University of Cambridge) for cell lines and the University of Oxford Department of Biomedical Services (BMS) Functional Genetics and JR Hospital BMS facilities for technical support.

## Funding

This work was funded by a Cancer Research UK (CRUK) Senior Cancer Research Fellowship RCCSCF-Nov21\100004 (awarded to J.R.C.), which provides salary and research support to J.R.C. and L.S. A.K. was previously supported by EMBO Long-Term Fellowship (EMBO - ALT 542-2020). This research was funded by UKRI via Medical Research Council Molecular Hematology Unit program funding to J.R.C. (MC_UU_00016/19 and MC_UU_00029/2). J.R.C. is the recipient of Lister Institute Research Prize Fellowship funding.

## Author contributions

Conceptualization: LS, JRC

Methodology: LS, AK, JRC

Investigation: LS

Formal analysis: LS, JRC

Supervision: LS, JRC

Writing—original draft, review and editing: LS, JRC

Funding acquisition: JRC

## Competing interests

Authors declare that they have no competing interests.

## Data and materials availability

All data are available in the main text or the extended data materials.

## Notes

### Competing Interest Statement

The authors have declared no competing interest.

