## Supplementary material for "ATM safeguards DNA replication at endogenous base lesions": Complete Supplementary Figures 1-8

**Affiliations:**

**Corresponding author:**

Supplementary Figures 1-10

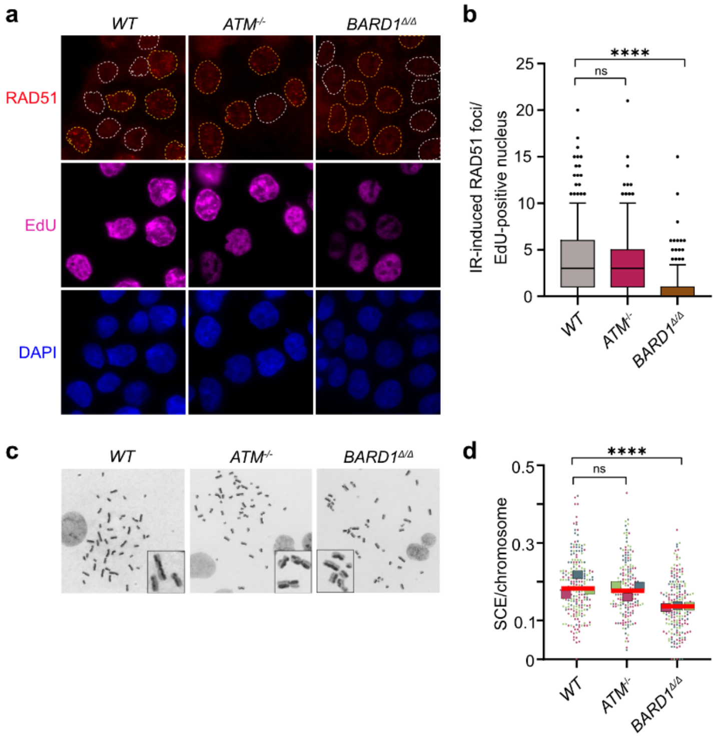


**Supplementary Figure 1**

**Figure S1. Homologous recombination (HR) activity is intact in *ATM*-deficient cells (related to Fig. 1)**

**a,b,** Immunofluorescence imaging of IR-induced RAD51 foci in indicated $BARD1^{AID/AID}$ HCT-116 cell lines following 2 h recovery from irradiation (5 Gy). Representative images **(a)**, EdU -positive and -negative nuclei are outlined in yellow and white, respectively. CellProfiler based quantification of RAD51 foci **(b)** across *n=3* biological experiments, each comprising >200 cells per condition. Horizontal bars, boxes, and whiskers denote median, 25–75, and 5–95 percentile values, respectively. Significance, two-sided Kruskal–Wallis H test with Dunn's correction for multiple comparisons. ****P ≤ 0.0001; ns non-significant.

**c,d,** Sister-chromatid exchanges (SCE) analysis in metaphase chromosome spreads of indicated cell lines cultured with 10 μM BrdU for two cell doublings. Representative images **(c)** and superplots **(d)** show average SCE frequency per chromosome across *n=3* biological experiments (>40 metaphases per experiment). Bars, median value. Colored squares, median SCE frequency from individual experiments. Significance, two-sided Kruskal–Wallis H test with Dunn's correction for multiple comparisons. ****P ≤ 0.0001; ns- non-significant.

Throughout the figure, *WT* denotes non-auxin-treated $BARD1^{AID/AID}$ cells. In the experiments involving $BARD1^{\Delta/\Delta}$ cells, all the indicated $BARD1^{AID/AID}$ cell lines were cultured in doxycycline (2 μg/ml) for 24 h before auxin (IAA, 1 mM) or dimethyl sulfoxide (DMSO) (carrier control).

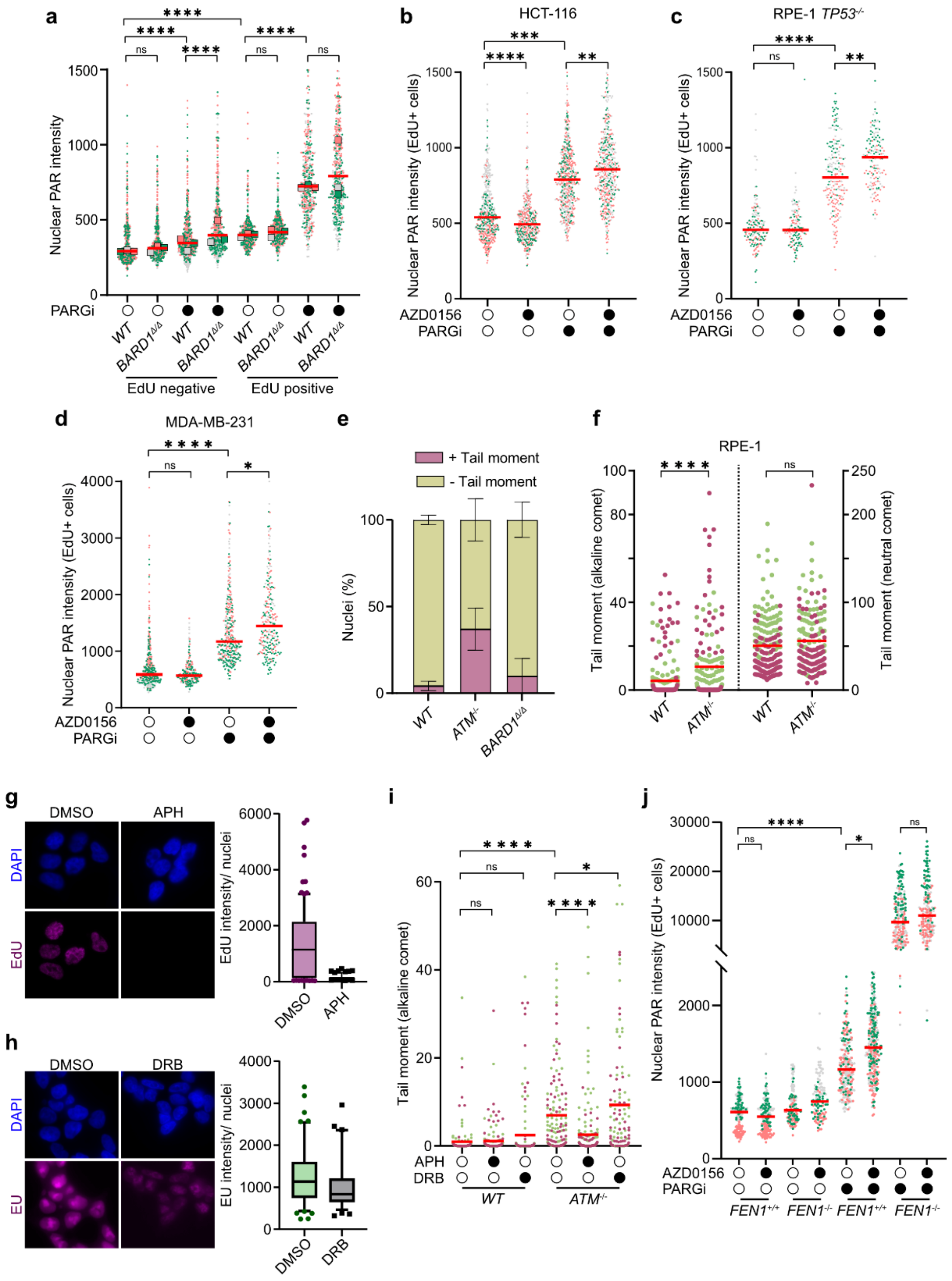


**Supplementary Figure 2**

**Figure S2. Accumulation of PARP-activating SSBs characterizes *ATM*-deficient cells (related to Fig. 1)**

**a,** Quantification of nuclear PAR levels in indicated *BARD1*$^{AID/AID}$ HCT-116 cell lines. Superplots show integrated nuclear PAR intensity across *n=3* biological experiments, each represented in different colors, with bars denoting median. Colored squares, median PAR signal from individual replicates. Significance, two-way ANOVA mixed model with repeat measures and Sidak's correction for multiple comparisons. ****P ≤ 0.0001; ***P ≤ 0.001; **P ≤ 0.01; ns- non-significant.

**b-d,** Quantification of nuclear PAR levels in EdU pulse-labelled *BARD1*$^{AID/AID}$ HCT-116 **(b),** *TP53*$^{-/-}$ RPE-1 cells **(c),** and MDA-MB-231 **(d)** cells treated with AZD0156 (500 nM), or carrier (DMSO, control) for 2 h. PARG inhibitor (PARGi) or carrier (mock-treatment) was added 45 min before fixation. Scatter plots, integrated intensity of PAR signals per EdU positive nucleus quantified across *n=3* biological experiments, each represented in a different color. Bars, median PAR signals. Colored squares, median PAR signal from individual experiments. Significance, two sided Kruskal-Wallis H test with Dunn's correction for multiple comparisons, ****P ≤ 0.0001; ***P ≤ 0.001; **P ≤ 0.01; *P ≤ 0.05; ns- non-significant.

**e,** Quantification of DNA damage in the indicated *BARD1*$^{AID/AID}$ HCT-116 cells by alkaline comet assay. Stacked bar charts show proportion of nuclei in **(Fig. 1g)** positive for an alkaline tail moment signal exceeding 0.5. Mean ± s.d. Scatter plots, comet tail moments across *n=3* biological experiments (80-100 cells per experiment, each color represents one replicate). Horizontal bars, median values. Significance, two sided Kruskal-Wallis H test with Dunn's correction for multiple comparisons. ****P ≤ 0.0001; ns, non-significant.

**f,** Quantification of DNA damage in the indicated RPE-1 cells lines by alkaline comet assay (left) and neutral comet assay (right). Scatter plots show comet tail moments across *n=2* biological experiments (80-100 cells per experiment, each color represents one replicate). Horizontal bars, median values. Significance, two sided Kruskal-Wallis H test with Dunn's correction for multiple comparisons. ****P ≤ 0.0001; ns, non-significant.

**g,h,** Immunofluorescence imaging of EdU and EU incorporation in *BARD1*$^{AID/AID}$ HCT-116 cells. Cells were treated with aphidicolin (APH; 1 µM; 1 h) to inhibit DNA replication, or with 5,6-dichlorobenzimidazole (DRB; 100 µM; 1 h) to inhibit DNA transcription. Representative images and quantification of integrated EdU **(g)** or EU **(h)** intensity quantification across *n=2* biological experiments (>150 cells per condition). Horizontal bars, boxes, and whiskers denote median, 25–75, and 5–95 percentile values, respectively.

**i,** Alkaline comet assays in the indicated *BARD1*$^{AID/AID}$ HCT-116 cell-lines from **g,h.** Scatter plots, tail moments across *n=2* biological experiments, (80-100 cells per experiment, each color represents one replicate). Horizontal bars, median values. Significance, two sided Kruskal-Wallis H test with Dunn's correction for multiple comparisons. ****P ≤ 0.0001; *P ≤ 0.05; ns, non-significant.

**j,** Quantification of nuclear PAR levels in S-phase staged RPE-1 cells incubated with or without AZD0156 (100 nM; 1 h) followed by PARGi (1 mM), or mock (DMSO) for 45min. Data, integrated PAR intensity values for EdU positive nuclei, across *n=2* biological experiments (each represented in a different color). Horizontal bars, median values. Significance, two sided Kruskal-

Wallis H test with Dunn's correction for multiple comparisons, ****P ≤ 0.0001, ***P ≤ 0.001; *P ≤ 0.05; ns, non-significant

*WT* refers to non-auxin-treated *BARD1*$^{AID/AID}$ cells. In the experiments involving *BARD1*$^{\Delta/\Delta}$ cells, all the indicated *BARD1*$^{AID/AID}$ cell lines were cultured in doxycycline (2 μg/ml) for 24 h before auxin (IAA, 1 mM) or dimethyl sulfoxide (DMSO) (carrier control).

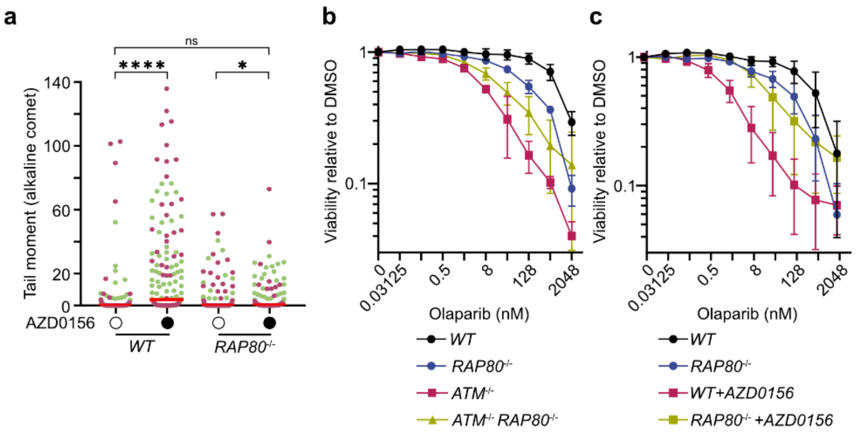


**Supplementary Figure 3**

**Figure S3. BRCA1-A-dependent SSBs accumulation drives PARPi toxicity in *ATM*-deficient cells (related to Fig. 2)**

**a,** Alkaline comet tail moment in indicated cell lines, data from *n=2* biological experiments (80-100 cells per experiment; each color denotes one replicate). Horizontal bars, median. Significance, two sided Kruskal-Wallis H test with Dunn's correction for multiple comparisons. ****P ≤ 0.0001; *P ≤ 0.05; ns, non-significant.

**b,c,** Resazurin survival assay of indicated non-auxin-treated $ATM^{-/-}$ cells **(b)** and $BARD1^{AID/AID}$ HCT-116 treated with ATMi (AZD0156, 500nM) or mock (DMSO) **(c)** added 1 h prior adding olaparib. *n=3* biological experiments, mean ± s.d.

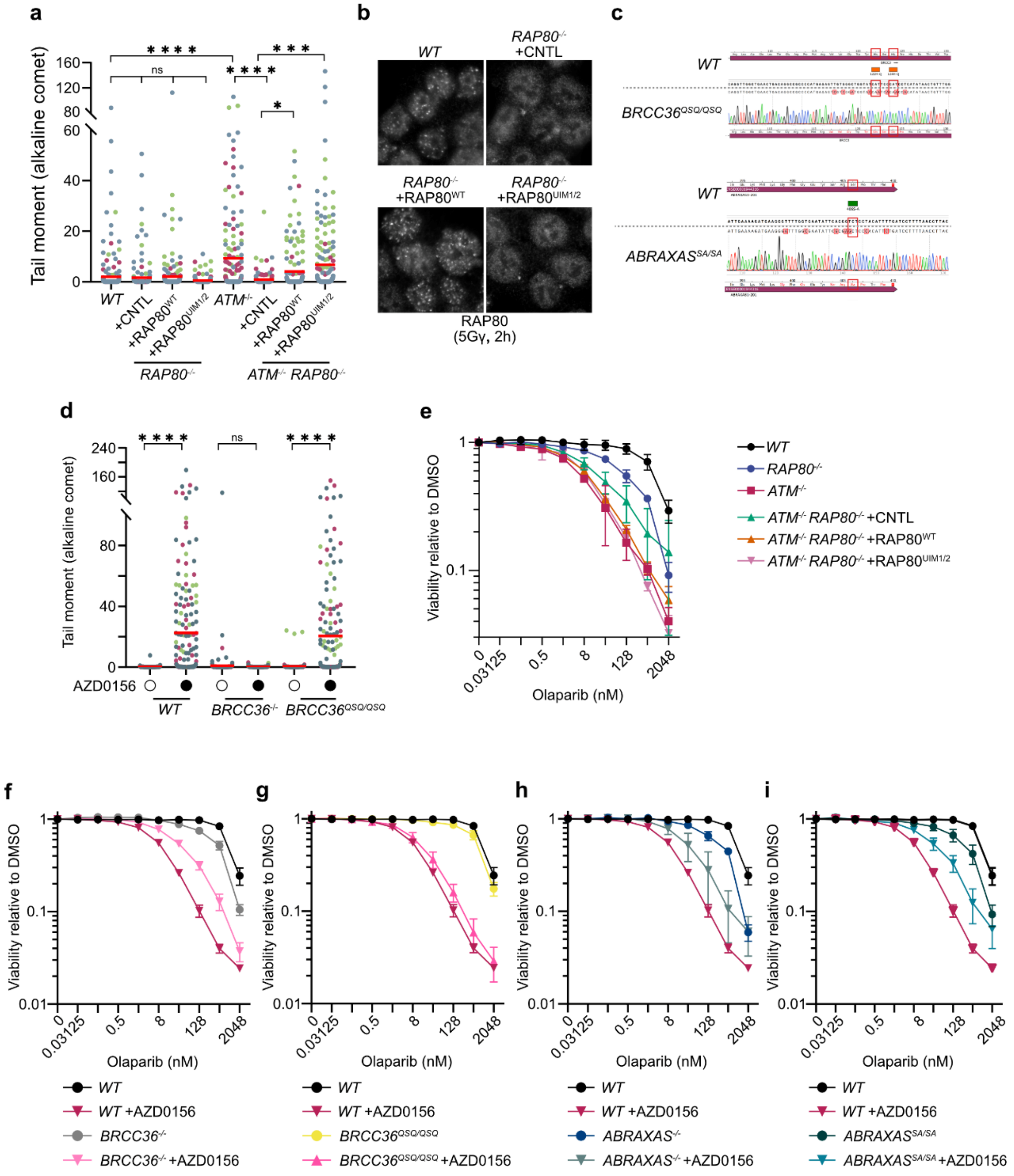


**Supplementary Figure 4**

**Figure S4. BRCA1 interaction with BRCA1-A complex is essential for SSBs accumulation driving PARPi toxicity in *ATM*-deficient cells (related to Fig. 2)**

**a,** Alkaline comet tail moment in indicated cell lines, data from *n=3* biological experiments (80-100 cells per experiment; each color denotes one replicate). Horizontal bars, median. Significance, two sided Kruskal-Wallis H test with Dunn's correction for multiple comparisons. ****$P \leq 0.0001$; ***$P \leq 0.001$; *$P \leq 0.05$; ns, non-significant.

**b,** Representative immunofluorescence images of IR-induced RAP80 foci in $BARD1^{AID/AID}$ HCT-116 cell lines complemented with wild type *RAP80*, $RAP80^{UIM1/2}$, or control (GST) expression transgenes following 2 h recovery from irradiation (5 Gy).

**c,** Sanger sequencing traces confirming the genotype of $BRCC36^{QSQ/QSQ}$ and $ABRAXAS^{S406A/S406A}$ cell lines.

**d,** Alkaline comet tail moment in indicated cell lines, data from *n=3* biological experiments (80-100 cells per experiment; each color denotes one replicate). Horizontal bars, median. Significance, two sided Kruskal-Wallis H test with Dunn's correction for multiple comparisons. ****$P \leq 0.0001$; *$P \leq 0.05$; ns, non-significant.

**e,** Survival assay of indicated non-auxin-treated $ATM^{-/-}$ $RAP80^{-/-}$ $BARD1^{AID/AID}$ HCT-116 complemented with $RAP80^{WT}$, $RAP80^{UIM1/2}$, or control (GST) expression transgenes. Resazurin cell viability assays, *n=3* biological experiments, mean ± s.d.

**f-i,** Survival assay of indicated non-auxin-treated $BARD1^{AID/AID}$ HCT-116 treated with ATMi (AZD0156, 500nM) or mock (DMSO) added 1 h prior adding olaparib. Resazurin cell viability assays, *n=3* biological experiments, mean ± s.d.

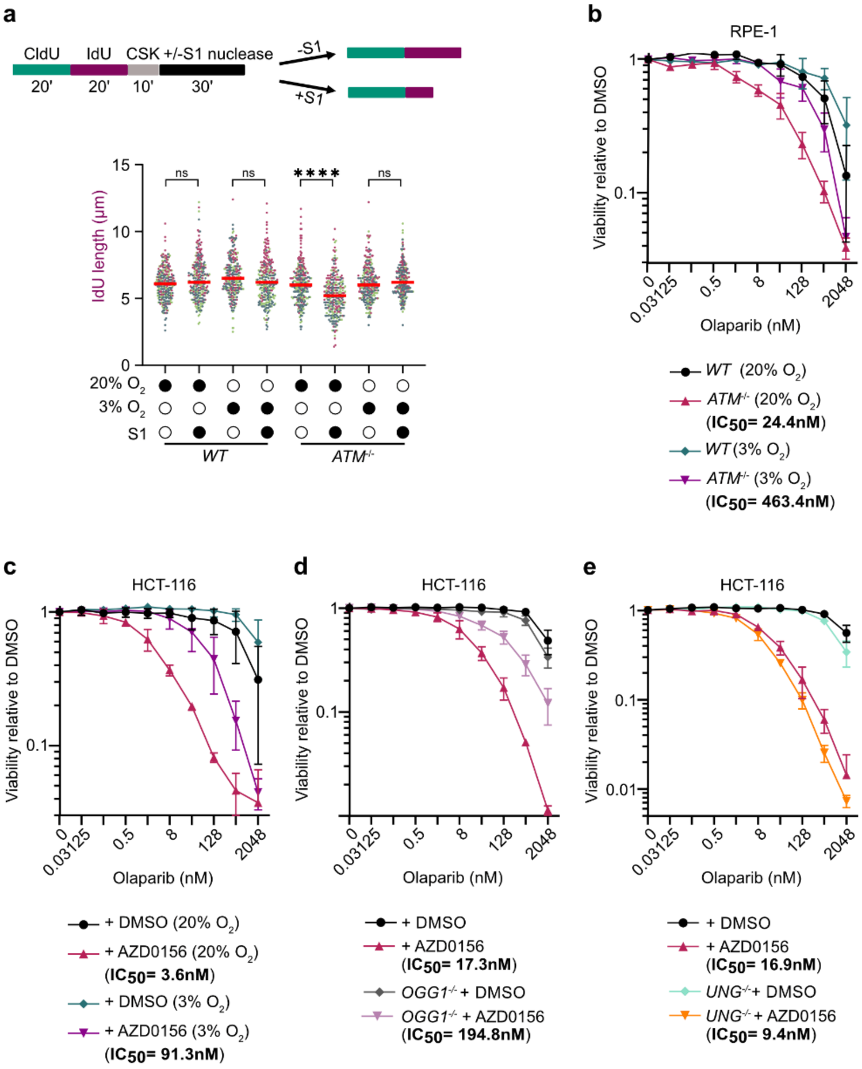


**Supplementary Figure 5**

**Figure S5. Base-excision repair intermediates drive PARPi sensitivity in *ATM*-deficient cells (related to Fig. 4)**

**a,** Schematic of nascent-strand ssDNA gap detection by S1-nuclease coupled DNA fiber assay. IdU, iododeoxyuridine; CldU, chlorodeoxyuridine (top panel). Scatter plot (bottom panel) showing IdU tract lengths in DNA fibers from the indicated $BARD1^{AID/AID}$ HCT-116 cells lines cultivated under ambient (20%) or physiological (3%) $O_2$ condition. Horizontal bars, median values of >250 fibers per cell line in n=3 biological experiments, each represented in a different color. Significance was determined by two sided Kruskal-Wallis H test with Dunn's correction for multiple comparisons. ****P ≤ 0.0001; ns, non-significant.

**b-e,** Survival assay of indicated RPE-1 cell-lines **(b,** *WT* referred to parental $ATM^{+/+}$ cells**)** and indicated non-auxin-treated $BARD1^{AID/AID}$ HCT-116 treated with ATMi (AZD0156, 500nM) or mock (DMSO) **(c-e)**. Cells were cultivated under ambient (20%) and physiological (3%) $O_2$ and exposed to the presence of indicated doses of olaparib. ATM inhibitor (AZD0156; 500 nM) was added 1 h prior adding olaparib. Resazurin cell viability assays, *n=3* biological experiments, mean ± s.d.

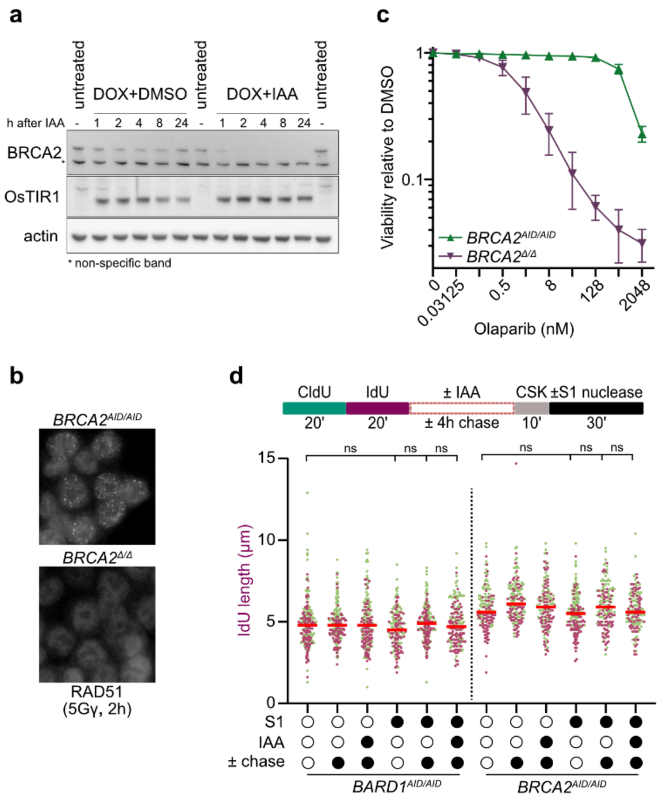


Supplementary Figure 6

**Figure S6. Homologous recombination (HR) mediates nascent-strand gap repair behind replication forks in *ATM*-deficient cells (related to Fig. 6)**

**a,** Immunoblot analysis of $BRCA2^{AID/AID}$ HCT-116 cell extracts at the indicated time points after addition of doxycycline (DOX) to induce OsTIR1 expression and auxin (IAA) to deplete BRCA2 protein. Representative of *n=2* independent experiments. *Indicates a non-specific band.

**b,** Representative immunofluorescence images of IR-induced RAD51 foci in $BRCA2^{AID/AID}$ HCT-116 cell lines treated with auxin (IAA, $BRCA2^{\Delta/\Delta}$) following 2 h recovery from irradiation (5 Gy).

**c,** Survival assay of $BRCA2^{AID/AID}$ HCT-116 treated with auxin (IAA) or mock (DMSO) added 1 h prior adding olaparib. Resazurin cell viability assays, *n=3* biological experiments, mean ± s.d.

**d,** Nascent-strand gap detection by S1-nuclease coupled DNA fiber assay. Schematic (top panel) shows, following IdU labelling, cells were allowed to recover in the complete media supplemented with auxin (IAA) or DMSO for 4 h prior S1 nuclease treatment. IdU, iododeoxyuridine; CldU, chlorodeoxyuridine. Scatter plots (bottom panel) showing IdU tract lengths in DNA fibers from the indicated $BARD1^{AID/AID}$ or $BRCA2^{AID/AID}$ HCT-116 cells lines. Horizontal bars, median values of >250 fibers per cell line in *n=2* biological experiments, each represented in a different color. Significance was determined by two sided Kruskal-Wallis H test with Dunn's correction for multiple comparisons. ns, non-significant.

In experiments involving $BARD1^{\Delta/\Delta}$ or $BRCA2^{\Delta/\Delta}$ cells, all the indicated $BARD1^{AID/AID}$ or $BRCA2^{AID/AID}$ cell lines were cultured in doxycycline (2 µg/ml) for 24 h before auxin (IAA, 1 mM) or dimethyl sulfoxide (DMSO) (carrier control).

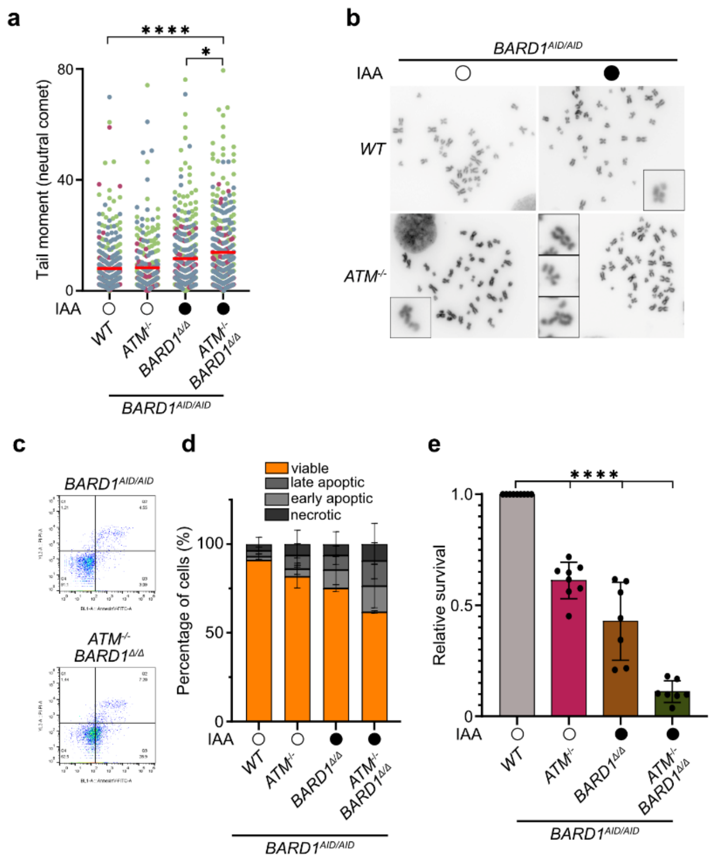


**Supplementary Figure 7**

**Figure S7.** ***ATM*****-deficient cells are addicted to homologous recombination (HR) (related to Fig. 6)**

**a,** Neutral comet assays of indicated *BARD1*$^{AID/AID}$ HCT-116 cell lines. Scatter plots, alkaline/neutral comet tail moments from *n=3* biological experiment (80-100 cells per experiment, each color represents one replicate). Significance, two sided Kruskal-Wallis H test with Dunn's correction for multiple comparisons. ****P ≤ 0.0001; ns, non-significant.

**b,** Representative metaphase images from *n = 3* biological experiments using the indicated *BARD1*$^{AID/AID}$ HCT-116 cell lines.

**c,d,** Flow cytometry analysis apoptotic markers (AnnexinV and propidium iodide) in indicated *BARD1*$^{AID/AID}$ HCT-116 cell-lines 96 h after treatment with auxin (IAA) or carrier (DMSO). Gating strategy **(c)** and quantification **(d)**. Data, *n=3* biological experiments, mean ± s.d.

**e,** Quantification of 7-day clonogenic cell survival of the indicated *BARD1*$^{AID/AID}$ cell lines. Data, mean ± s.d of *n=3* biological experiment (each with >2 technical repeats). Significance, ordinary one-way ANOVA. ****P ≤ 0.0001.

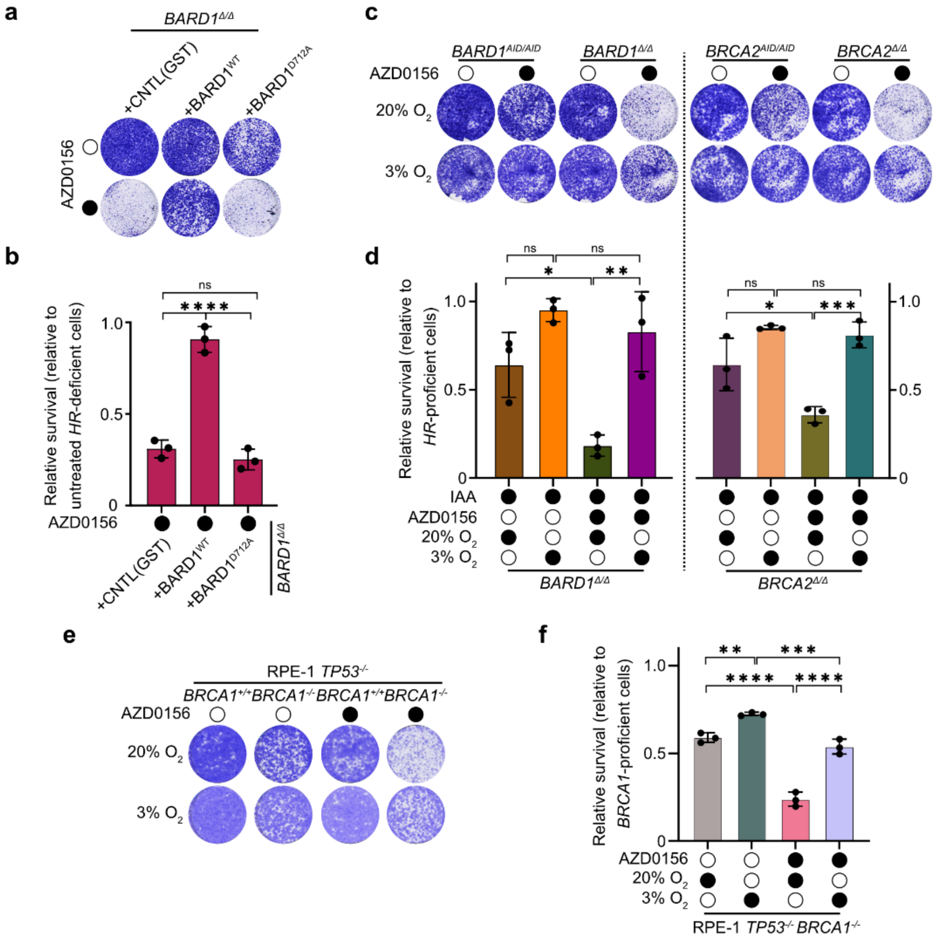


**Supplementary Figure 8**

**Figure S8. Nascent-strand SSBs drive homologous recombination (HR) addiction in *ATM*-deficient cells (related to Fig. 6)**

**a,b,** Seven-day clonogenic survival assays with the indicated $BARD1^{AID/AID}$ HCT-116 cells, complemented with wild type *BARD1,* $BARD1^{D712A}$, or control (GST) expression transgenes. Representative images **(a)** and quantification **(b)** of crystal violet staining. Data, mean ± s.d. of *n=3* biological experiments (each with >2 technical repeats). Significance was determined by ordinary one-way ANOVA with Tukey's correction for multiple comparisons. ****P ≤ 0.0001; ns, non-significant.

**c-f,** Seven-day clonogenic survival assays with the indicated $BARD1^{AID/AID}$ and $BRCA2^{AID/AID}$ cell lines **(c,d)**. $BARD1^{AID/AID}$, $BRCA2^{AID/AID}$ cells treated with ATM inhibitor (AZD0156, 500 nM) **(c,d)** and $BRCA1^{-/-}$ $TP53^{-/-}$ RPE-1 cells **(e,f)** treated with ATM inhibitor (AZD0156, 100 nM) or mock (DMSO) grown at ambient (20%) or physiological (3%) $O_2$ conditions. Representative images **(c,e)** and quantification **(d,f)** of crystal violet staining. Data, mean ± s.d. of *n=3* biological experiments (each with >2 technical repeats). Significance was determined by ordinary one-way ANOVA with Tukey's correction for multiple comparisons. ****P ≤ 0.0001; ***P ≤ 0.001; **P ≤ 0.01; * P ≤ 0.05; ns, non-significant.

In experiments involving $BARD1^{\Delta/\Delta}$ and $BRCA2^{\Delta/\Delta}$ cells, all the indicated $BARD1^{AID/AID}$ $BRCA2^{AID/AID}$ cell lines were cultured in doxycycline (2 µg/ml) for 24 h before auxin (IAA, 1 mM) or dimethyl sulfoxide (DMSO) (carrier control).
